# Mechano-regulation of GLP-1 production by Piezo1 in intestinal L cells

**DOI:** 10.1101/2024.04.01.587569

**Authors:** Yanling Huang, Haocong Mo, Jie Yang, Luyang Gao, Tian Tao, Qing Shu, Wenying Guo, Yawen Zhao, Jingya Lyu, Qimeng Wang, Jinghui Guo, Hening Zhai, Linyan Zhu, Hui Chen, Geyang Xu

## Abstract

Glucagon-like peptide 1 (GLP-1) is a gut-derived hormone secreted by intestinal L cells and vital for postprandial glycemic control. As open-type enteroendocrine cells, whether L cells can sense mechanical stimuli caused by chyme and thus regulate GLP-1 synthesis and secretion is unexplored. Our study showed expression of Piezo1 in intestinal L cells. Its level varied in different energy status and correlates with blood glucose and GLP-1 levels. Mice with L cell-specific loss of Piezo1 (*IntL-Piezo1^-/-^*) exhibited impaired glucose tolerance, increased body weight, reduced GLP-1 production and decreased CaMKKβ/CaMKIV-mTORC1 signaling pathway under normal chow diet or high fed diet. Activation of the intestinal Piezo1 by its agonist Yoda1 or intestinal bead implantation increased the synthesis and secretion of GLP-1, thus alleviated glucose intolerance in diet-induced-diabetic mice. Overexpression of Piezo1, Yoda1 treatment or stretching stimulated GLP-1 production and CaMKKβ/CaMKIV-mTORC1 signaling pathway, which could be abolished by knockdown or blockage of Piezo1 in primary cultured mouse L cells and STC-1 cells. These findings suggest a previously undiscovered mechano-regulation of GLP-1 production in L cells, which may shed new light on the treatments of diabetes.

## Introduction

The gastrointestinal (GI) tract represents the largest endocrine organ in the human body. The enteroendocrine cells (EECs) located throughout the GI tract secrete a large number of gastrointestinal hormones to regulate a variety of physiological processes and are key regulators for energy homeostasis (Bany Bakar et al., 2023). GLP-1 is one of the gut-derived peptide hormones essential for postprandial glycemic control (Song et al., 2019). It is produced from Proglucagon (Gcg) by proprotein convertase in the intestinal L cells, a group EECs predominantly situated in the distal gut (Drucker, 2006; Rouillé et al., 1997). The circulating GLP-1 levels rapidly increase after meal and reduce postprandial blood glucose fluctuations by augmenting insulin secretion, suppressing glucagon secretion and slowing gastric emptying (Drucker, 2006; Willms et al., 1996). Nowadays, GLP-1-based therapy is well-recognized and commonly used in treatment of Type 2 Diabetes Mellitus (T2DM) (Saxena et al., 2021; Tan et al., 2022). Elucidation of the mechanism that regulates GLP-1 production is essential for the development of new drug targets for the treatment of diabetes.

EECs can be divided into two categories according to its morphology: open type and closed type. The open type EECs possess microvilli protruding into the gut lumen and have direct contact with the luminal contents. In contrast, the closed type EECs are located basolaterally without direct contact with the lumen (Gribble & Reimann, 2016). Both types of EECs synthesize and store peptides or hormones in secretory granules and release them by exocytosis at the basolateral membrane upon mechanical, chemical or neural stimulation (Atanga et al., 2023). As open-type EECs, L cells received both chemical and mechanical signals from the luminal contents, and neural signals from the nerves (Furness et al., 2013). It has been well-documented that nutrients such as glucose, lipids, and amino acids in the intestinal lumen can stimulate the secretion of GLP-1 from L cells (Diakogiannaki et al., 2012). GLP-1 secretion can also be stimulated by intrinsic cholinergic nerves (Anini et al., 2002; Drucker, 2006). However, whether and how L cells coordinate mechanical stimuli from intestinal lumen to regulate GLP-1 production remain poorly understood.

Piezo channels, including Piezo1 and Piezo2 have recently been identified as mechanosensitive ion channels involved in the sensation of multiple mechanical stimuli, such as shear stress, pressure, and stretch (Gudipaty et al., 2017; Li et al., 2014; Romac et al., 2018). They allow the influx of cations such as Ca^2+^ and Na^+^ in response to mechanical tension and converts mechanical stimuli into various electrical and chemical signals. Piezo1 plays a crucial role in blood pressure regulation, red blood cell volume regulation, bone homeostasis, pulmonary and cardiac functions (Cahalan et al., 2015; Lai et al., 2022; Wang et al., 2023; Wang et al., 2016). Previous studies have reported that Piezo1 is expressed in the intestinal epithelium, regulating gut peristalsis, barrier function, mucus secretion, and inflammation (Jiang et al., 2021; Liu et al., 2022; Sugisawa et al., 2020; Xu et al., 2021). Interestingly, accumulating evidence demonstrates the regulation of insulin and ghrelin secretion by Piezo1 (Deivasikamani et al., 2019; Ye et al., 2022; Zhao et al., 2024). Recent studies have also reported that Piezo2 is expressed in a population of EECs and convert force into serotonin release (Alcaino et al., 2018; Treichel et al., 2022). These findings suggest a critical role of Piezo channels in the mechano-regulation of hormone production. However, whether Piezo channels are expressed L cells and play a role in GLP-1 production remain unknown.

Here, we present evidence that Piezo1 channels on intestinal L cells mediate the mechanosensing from intestinal contents and trigger the synthesis and secretion of GLP-1 through CaMKKβ/CaMKIV-mTORC1 signaling pathway, thus regulating glucose homeostasis. Our research provides new insights for the treatment of T2DM and a new theoretical basis for hypoglycemic medicines targeting Piezo1.

## Methods

### Collection of Human Intestine Samples

Male obese participants with type 2 diabetes (n=6, BMI=45.87±4.889 kg/m^2^) and one-year post-RYGB patients (n=6, BMI=25.48±1.085 kg/m^2^) were recruited in current study. Written informed consent was obtained from each donor. The study protocol was approved by the Institutional Review Board of Jinan University. Mucosal biopsies were obtained from human intestines by using a colonoscopy (CF-HQ290I; Olympus).

### Genetic Mouse Generation

#### Villin-Flippase (Vil-Flp) mice

*Vil-Flp* knock-in mouse model was developed by Shanghai Model Organisms Center, Inc. The targeting construct was designed to insert a 2A-Flp-WPRE-pA coexpression cassette into the stop codon of mouse *Vil1* gene via homologous recombination using CRISPR/Cas9 system. 5’-AGCCCCTACCCTGCCTTCAA-3’ was chosen as Cas9 targeted guide RNA (sgRNA). The donor vector, sgRNA and Cas9 mRNA was microinjected into C57BL/6J fertilized eggs. F0 generation mice positive for homologous recombination were identified by long PCR. The primers (I-IV) used for detection of the correct homology recombination were I: 5’-ACTTCAGGCCTAACGCTCAC-3’ and II: 5’-TGTCCTGCAGGCAGAGAAAG-3’ for the correct 5’ homology arm recombination, and III: 5’-GTGCCGTCTCTAAGCACAGT-3’and IV:5’-TGTTGGTGCTTCGGAGTGTT-3’for the correct 3’ homology arm recombination. The PCR products were further confirmed by sequencing. F0 mice were crossed with C57BL/6J mice to obtain *Vil-Flp* heterozygous mice.

#### Flp-dependent Glucagon-Cre (FGC) mice

*FGC* mouse model was developed by Shanghai Model Organisms Center, Inc. The targeting construct was designed to insert an IRES-F3-Frt-Wpre-pA-Cre-Frt-F3 expression cassette into the 3’ UTR of mouse *Gcg* gene of via homologous recombination using CRISPR/Cas9 system. 5’-ATGCAAAGCAATATAGCTTC-3’ was chosen as Cas9 targeted guide RNA (sgRNA). The donor vector, sgRNA and Cas9 mRNA was microinjected into C57BL/6J fertilized eggs. F0 generation mice positive for homologous recombination were identified by long PCR. The primers (I-IV) used for detection of the correct homology recombination were I: 5’-TGCTACACAGGAGGTCTGTC-3’ and II: 5’-AGGCATGCTCTGCTATCACG-3’ for the correct 5’ homology arm recombination, and III: 5’-CCCTCCTAGTCCCTTCTCAGT-3’ and IV: 5’-GCCAAGGACATCTTCAGCGA-3’ for the correct 3’ homology arm recombination. The PCR products were further confirmed by sequencing. F0 mice were crossed with C57BL/6J mice to obtain *FGC* heterozygous mice.

#### IntL-Cre mice

*Vil-Flp* mice were crossed with *FGC* mice to obtain Intestinal L cell-specific Cre (*IntL-Cre*) mice.

#### IntL-Piezo1^-/-^ mice

*Piezo1^loxp/loxp^*mice (B6.Cg-Piezo1^tm2.1Apat^/J) purchased from Jackson laboratory were crossed with IntL-Cre mice to generate *IntL-Piezo1^-/-^*mice.

PCR is used to identify the genotype of mice during the subsequent mating and breeding process. The primers required for mouse genotyping are shown in the Supplementary information, Table S1.

### Mouse Validation

*mT/mG* reporter mice were purchased from Shanghai Model Organisms Center, Inc. *IntL-Cre* mice were bred with *mT/mG* reporter mice to further validate Cre recombinase activity and specificity. Every single *IntL-Cre* mouse was confirmed by *mT/mG* reporter mice before breeding with *Piezo1^loxp/loxp^* mice to generate Intestinal L cell-Piezo1^-/-^ (*IntL-Piezo1^-/-^*) mice.

### Frozen Tissue Confocal Imaging

*IntL-Cre-mT/mG* reporter mice were sacrificed. Fresh ileum and pancreas tissues were collected and embedded in O.C.T. compound for histological analysis immediately. Slices of the tissues were cut for confocal imaging, which was protected from light. Fluorescence was detected by laser scanning confocal microscopy (Li et al., 2022).

### Animal Housing and Treatment

Male mice were maintained on a 12-hour light/12-hour dark cycle environment. Normal chow diet (NCD) or a high-fat diet (HFD) and water were available ad libitum unless specified otherwise. The animal protocols were approved by the Animal Care and Use Committee of Jinan University.

Male *IntL-Piezo1^-/-^*mice and age-matched control littermates (*Piezo1^loxp/loxp^*,

*Vil-Flp*, *FGC*, *IntL-Cre* mice) fed with NCD or HFD were used in the experiments.

Male *Piezo1^loxp/loxp^*and *IntL-Piezo1^-/-^* mice fed with 10 week-high fat diet were intraperitoneally injected with normal saline (NS) or the GLP-1R agonist Exendin-4 (100 ug/kg body weight) for 7 consecutive days.

High fat diet induced diabetic mice were randomly divided into 3 groups. When indicated, animals were injected intraperitoneally with Vehicle, Yoda1 (2 μg per mouse) or GsMTx4 (250 μg/kg) plus Yoda1 for 7 consecutive days.

High fat diet treated *IntL-Piezo1^-/-^* mice were randomly divided into 2 groups. When indicated, animals were injected intraperitoneally with Vehicle, Yoda1 (2 μg per mouse) for 7 consecutive days.

Diet induce diabetic C57BL/6J mice were divided into sham and intestinal bead implantation groups.

### Food and water intake detection

The food and water intake were quantified using metabolic cages (Cat 41853, Ugo Basile, Comerio, Italy). The mice were individually housed in these specialized cages and given a period of 3 days to acclimate before data collection began. They had unrestricted access to food and water, which was continuously monitored throughout the study. The 41850-010 software/interface package, consisting of EXPEDATA (for data analysis) and METASCREEN (for data collection) software, along with the IM-2 interface module, was employed to record and analyze the data.

### Intraperitoneal Glucose Tolerance Test

Mice were fasted for 12 hours before measuring their fasting glucose levels. An intraperitoneal glucose tolerance test (IPGTT) was performed by administering 1.5 g/kg body weight of glucose. Blood glucose concentrations were measured at specified time points using a glucometer by collecting tail vein blood samples.

### Insulin Tolerance Test

Mice were subjected to a 4-hour fast before measurement of fasting glucose were taken. Insulin tolerance tests (ITT) were conducted with a dose of 0.75 U/kg body weight of insulin. Blood glucose levels were measured at specified time points.

### Intestinal Bead implantation

High-fat diet-induced type 2 diabetic C57BL/6J mice were fasted 6 to 8 h before the operation. Standard aseptic procedures were used throughout the operation. Intestinal bead implantation was similar to gastric bead implantation described in our previous study (Zhao et al., 2024). Briefly, a 1cm incision was made on the abdominal wall to expose the intestine. A 1 cm incision was made approximately 1cm above the ileocecal region. A 2.5 mm diameter bead was implanted into the ileum of the mouse through an incision. Then the wound was closed with suture. Finally, the abdominal wall was closed with suture. For sham operation, all the procedures were the same as the bead implantation except that the bead was not implanted.

### Stretching of isolated ileum

About 2cm ileum was isolated from control and *IntL-Piezo1^-/-^* mice and kept in the specimen chamber filled with Tyrode’s solution (KCl 0.2g/L, NaCl 8g/L, CaCl_2_ 0.2g/L, MgCl_2_ 0.1g/L, NaHCO_3_ 1g/L, NaH_2_PO_4_ 0.05g/L, Glucose 1g/L) of 37[ gassed with oxygen. The specimen was connected to the force transducer of organ bath system (HW200S, Techman, Chengdu, CN). Adjust the transducer to apply traction force of 1.5 grams on the tissue and maintained for two hours.

### Measurement of GLP-1 Secretion

The measurement of GLP-1 secretion was carried out according to previously described methods (Zhai et al., 2018). Samples were collected in the presence of aprotinin (2 μg/mL), EDTA (1 mg/mL) and diprotin A (0.1 mmol/L), and stored at −80 °C before use. GLP-1 levels were assayed using enzyme immunoassay kits following the manufacturer’s instructions.

### Histological Analysis

Tissues were collected, fixed with 4% paraformaldehyde, embedded in paraffin, and cut into 4μm sections. Standard protocols were followed for staining the sections with hematoxylin-eosin. Photomicrographs were captured under an inverted microscope (Leica, Germany).

### Immunofluorescence

Paraffin sections were dewaxed and rehydrated. After antigen retrieval in citrate buffer (pH6.0), sections were blocked with normal serum and then incubated with rabbit anti-Piezo1 (1:400) and mouse anti-GLP-1 (1:500) antibodies at 4°C overnight. The sections were then incubated with a mixture of secondary antibodies. Images were taken by laser scanning confocal microscopy (Leica SP8). Fluorescence intensity was quantified by ImageJ software.

### In situ hybridization

Paraffin sections were dewaxed and rehydrated. After antigen retrieval in in citrate buffer (pH6.0), the sections were incubated with Proteinase K (5ug/ml) at 37°C for the 15 minutes. Then the sections were hybridized with the probes overnight in a temperature-controlled chamber at 40°C. The Piezo1 probe sequences were as follows: 5′-CTGCAGGTGGTTCTGGATATAGCCC-3′,5′-AAGAAGCAGATCTCCAGCCCGAAT-3′, 5′-GCCATGGATAGTCAATGCACAGTGC-3′. After washing with SSC buffers, the sections were hybridized in pre-warmed branch probes at 40°C for 45 minutes. After washing with SSC buffers, the sections were hybridized with signal probe at 42°C for 3 hours. After washing with SSC buffers, the sections were blocked with normal serum and then incubated with mouse anti-GLP-1 (1:500) antibody at 4°C overnight followed by secondary antibody. Images were taken laser scanning confocal microscopy and the fluorescence signals were quantified by ImageJ.

### Western Blot Analysis

Tissues and cells were harvested. Ileal mucosa was scraped for protein extraction. Protein extraction was performed by using RIPA lysis buffer (50mM Tris PH 7.4, 150mM NaCl, 1% Triton X-100, 0.1% SDS, 1% Sodium deoxycholate, 1mM PMSF and protease inhibitor cocktail.), then 40 µg of proteins were loaded onto an SDS-PAGE gel for separation. After the separation, the proteins were transferred onto a nitrocellulose membrane. The membrane was then incubated in blocking buffer at room temperature for 1 hour. For overnight incubation, the membrane and primary antibody (at the recommended dilution as stated in the product datasheet) were immersed in primary antibody dilution buffer, with gentle agitation, at 4°C. Subsequently, the membrane was incubated with a secondary antibody that specifically recognizes and binds to the primary antibody. Finally, Western blotting luminol reagent was used to visualize bands. The grey scale values of the bands were measured using ImageJ software.

### RNA extraction, quantitative real-time PCR

RNA was extracted and reverse-transcribed into cDNAs using RT-PCR kit. Real-time PCR was performed as previously described (Zhai et al., 2018). Sequences for the primer pairs used in this study were shown in Supplementary information, Table S2.

### Isolation of Mouse Intestinal L Cells

A 5∼6cm long ileum segment was collected from the *IntL-cre-mT/mG* mouse. The tissue was washed with ice-cold PBS twice to remove the chyme in the lumen. The tissue was minced into 0.5mm^3^ pieces in ice-cold PBS and then digested in 100mIU collagenase I and 0.01g/mL trypsin at 37°C for 30min with rotation. After digested tissue was passed through 40μm and 30μm cell strainers sequentially, then centrifuged for 7min at 4°C. The cell pellet was resuspended in red cell lysis buffer and incubated for 10min at room temperature. The unlysed cells were collected by centrifugation and resuspended with 1mL cold PBS. The GFP positive cells was sorted by fluorescence-activated cell sorting (FASC) on Beckman Coulter MoFlo XDP cell sorter system.

### Cell Culture and Treatments

STC-1 cells were maintained in DMEM medium supplemented with 2.5% fetal bovine serum and 10% equine serum at 37 °C with 5% CO_2_ air. L cells were maintained in DMEM medium supplemented with 10% fetal bovine serum.

For cell transfection, cells were plated at optimal densities and grown for 48 h. Cells were then transfected with GFP, Piezo1-GFP, CaMKKβ, CaMKIV by using lipofectamine reagent according to the manufacturer’s instructions.

For stable knockdown of Piezo1 in STC-1 cells, short hairpin RNA (shRNA) sequences for mouse Piezo1 interference were cloned in to pLKO.1 vector. To produce lentivirus, psPAX2, pMD2G and pLKO.1 or pLKO.1-shPiezo1 plasmids (siPiezo1: CCAACCTTATCAGTGACTT) were co-transfected into 293T cells with lipofectamine 2000 reagent. Supernatant containing lentivirus was collected 48 hours after transfection and filtered through 0.45μm filter. The virus-containing supernatant was used to infect STC-1 cells. Forty-eight hours after infection, the STC-1 cells were subjected to 1ug/mL puromycin selection for 2-3 days.

For cell stretching, cells were grown in silicone elastic chambers coated by 0.1 % gelatin solution. After incubated at 37 °C for 24-48 hours, The chambers were subjected to mechanical stretch to 120% of their original length.

### Calcium Imaging

Cells were plated onto confocal dishes at optimal densities and grown for 24 h. Cells were loaded with the calcium fluorescent probe fluo-4 AM (1 μM) for 1 h at 37 °C, then the cells were treated with Yoda1 (5 μM) or GsMTx4. The intracellular calcium ions were measured at room temperature using a laser confocal microscope with an excitation wave length of 494 nm and an emission wave length of 516 nm. The change of fluorescent signal was presented as ΔF/F_0_ and plotted against time.

### Whole-cell Patch-Clamp recording

Borosilicate glass-made patch pipettes (BF150-86-7.5, Sutter Instrument Co, USA) were pulled with micropipette puller (P-1000, Sutter Instrument Co, USA) to a resistance of 3–5[MΩ after being filled with pipette solution: 138mM KCl, 10mM NaCl, 1mM MgCl_2_, 10mM Glucose and 10mM HEPES (pH 7.4). Cells were bathed in Margo-Ringer solution: 130mM NaCl, 5mM KCl, 1mM MgCl_2_, 2.5mM CaCl_2_, 10mM Glucose, 20mM HEPES (pH7.4). Whole-cell calcium currents of STC-1 cells were recorded with the EPC10 USB patch-clamp amplifier (HEKA, Germany) controlled by PatchMaster software.

### Key resource

Key reagents or resources are listed in the Supplementary information, Table S3.

### Statistical Analysis

All data were expressed as mean ± S.E.M. Statistical differences were evaluated by one-way ANOVA or Student’s t-test. The correlation was determined by Pearson analysis. P < 0.05 was considered significant. (*p < 0.05, **p < 0.01, *** p < 0.001, ns=not significance). In our study, the data collection and analysis processes were not conducted in a blinded manner with respect to the experimental conditions. For the administration of drugs to animals, we allocated mice of the same genetic background to various experimental cohorts using a randomization protocol. No data were excluded during the data analysis.

## Results

### Assessment of Piezo1 in human and mouse intestine in different energy status

*Piezo1* mRNA was found to be highly expressed in both mouse ileal mucosa and STC-1 cells (Figure1-figure supplement 1A). Moreover, Piezo1 was co-localized with GLP-1 in immunofluorescent staining on NCD fed mouse ileal sections, indicating its expression in L cells (Figure1-figure supplement 1B). Interestingly, increased body weight and impaired glucose tolerance were observed in high-fat diet-induced diabetic mice, while Piezo1 and Proglucagon expression levels in the ileal mucosa of diabetic mice were significantly lower than that in mice feed with normal chow diet (Figure1-figure supplement 1C-F). Moreover, ileal mucosal *Piezo1* mRNA levels were positively correlated with *Proglucagon* mRNA levels (Figure1-figure supplement 1G), but negatively correlated with the AUC of glucose tolerance test (Figure1-figure supplement 1H). Obese T2MD patients who underwent Roux-en-Y gastric bypass (RYGB) surgery showed decreased BMI (Figure1-figure supplement 1 I) and increased *Piezo1* and GLP-1 in ileal mucosa (Figure1-figure supplement 1J and K) compared to that before surgery. These findings indicated that Piezo1 is expressed in intestinal L cells and its level varies in different energy status.

### Generation and characterization of *IntL-Piezo1^-/-^* mice

To investigate the potential role of Piezo1 in GLP-1 production, we tried to knockout *Piezo1* in L cells by Cre-loxP system driven by an L cell-specific promoter. Proglucagon (encoded by *Gcg* gene) is mainly expressed in both L cells and pancreatic α cells (Jin, 2008). Villin-1 (encoded by *Vil1* gene) is expressed in gastrointestinal epithelium, including L cells, but not in pancreatic α cells (Maunoury et al., 1992; Rutlin et al., 2020). Since neither Gcg nor Villin are specific markers for L cells, we tried to generate a new line of mice enabling loss of Piezo1 expression specifically in the intestine L cell by combination of Flp-Frt and Cre-loxP system. We inserted a Flippase (Flp) expression cassette in the 3’UTR of *Vil1* to generate a *Vil1* promoter-driven Flp mice (*Vil-Flp*) (Figure 1A). Then, we generated Flippase-dependent Gcg promoter driven-Cre (*FGC*) mice by inserting an Frt-flanked Cre expression cassette in reverse orientation within the 3’- UTR of *Gcg* gene (Figure 1A). We further crossed the *Vil-Flp* mice with *FGC* mice to obtain L cell specific Cre mice (*IntL-Cre*), in which *Vil1* promoter-driven Flippase flipped the reverse Cre cassette into a correct orientation in Villin positive cells (including L cells, but not pancreatic α cells), and thus Cre can only be expressed under the *Gcg* promoter in L cells. The genotypes of the *Vil-Flp*, *FGC* and *IntL-Cre* mice were identified by PCR with specific primers (Figure 1B). The flipping of the reverse Cre cassette was validated by PCR, which confirmed that the flipping only occurred the intestine, but not in the pancreas (Figure 1C). To confirm the cell type specificity of Cre activity, we crossed *IntL-Cre* mice to *mT/mG* reporter mice. All tissues and cells of *mT/mG* mice express red fluorescence (membrane-targeted tdTomato; mT) at baseline, and switch to membrane-targeted EGFP in the presence of cell-specific Cre (Figure 1D). EGFP expression was only observed scatteredly in the intestine, but not in the pancreas, indicating the intestine-specific Cre activity in the *IntL-Cre* mice (Figure 1E). Finally, we bred *IntL-Cre* mice with *Piezo1^loxp/loxp^* mice to generate *IntL-Piezo1^-/-^* mice (Figure 1F).

**Figure 1:**
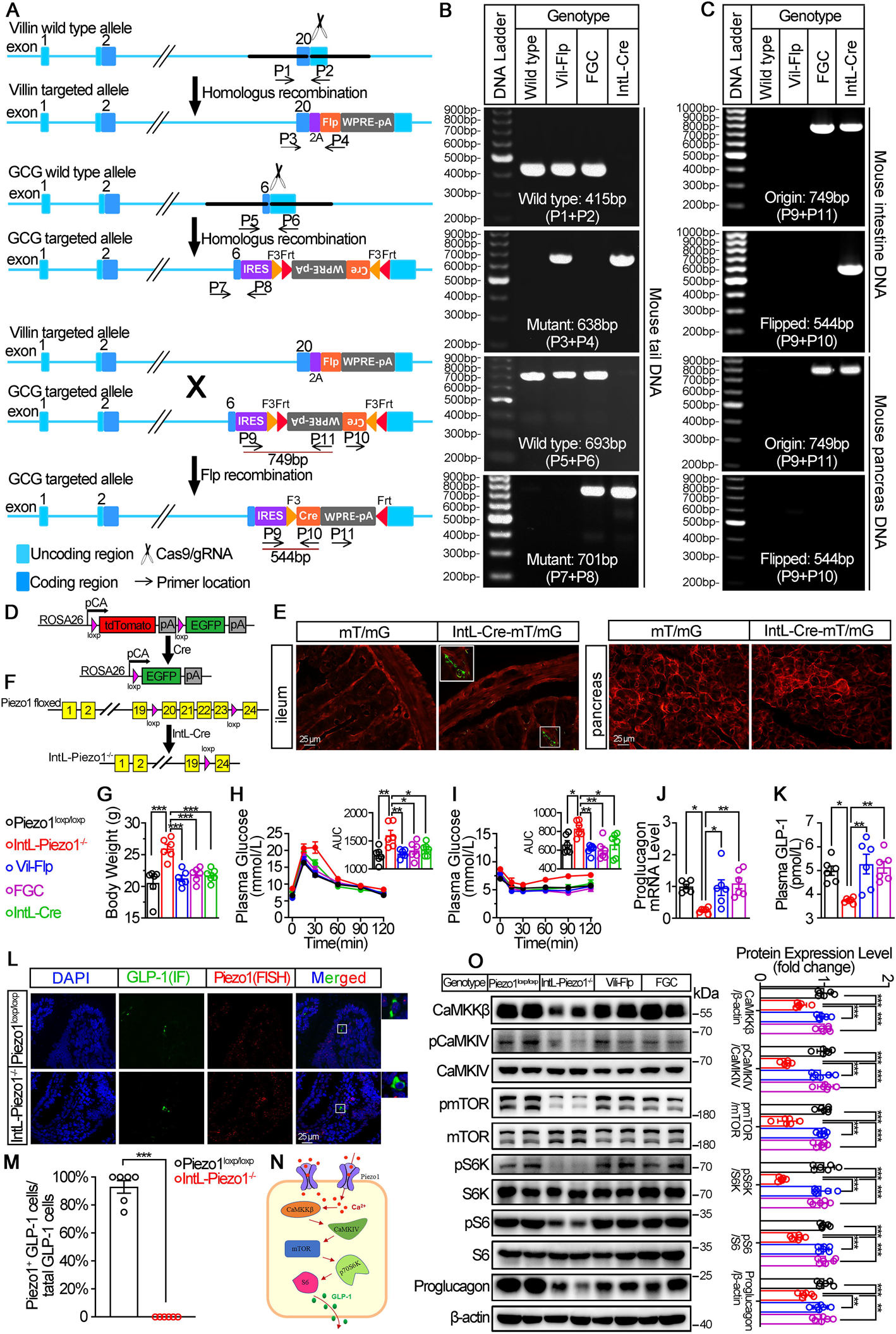
Generation, Validation and Characterization of *IntL-Piezo1^-/-^* mice. (A) Schematic description for the generation of Villin-Flippase (*Vil-Flp*) and Flippase-dependent Gcg-Cre (*FGC*) mice. Vil-Flp flip the inverted Cre gene in the Gcg-Cre cassette in *IntL-Cre* mice to restrict Cre expression in intestinal L cells. As shown, Locations of genotyping primers are also indicated. (B) Tail DNA genotyping PCR results using genotyping primer for *Vil-Flp*, *FGC* and Flippase-activated Cre (*IntL-Cre*) mice. (C) Intestine and pancreas DNA genotyping results. The “Original” band represents the original FGC cassette with inverted Cre, while the “Flipped” band represents recombined FGC cassette with Cre flipped into the correct direction. (D) Schematic description for the validation of *IntL-Cre* efficacy by crossing with *mT/mG* reporter mice. (E) Fluorescence was detected in the ileal and pancreatic tissues from *mT/mG* and *IntL-Cre-mT/mG* mice by frozen tissue confocal microscopy. Green fluorescence represents successful deletion of TdTomato and reactivation of EGFP in the Cre-expressing cells. (F) Schematic description for the generation of Intestinal L cell-Piezo1^-/-^ mice (*IntL-Piezo1^-/-^*) by crossing *Piezo1^loxp/loxp^* mice with *IntL-Cre* mice. (G) Body weight of 14- to 16-week-old male mice of the indicated genotypes fed NCD (n=6/group) (**H-I**) IPGTT (**H**) and ITT (**I**) and associated area under the curve (AUC) values of 14- to 16-week-old male mice of the indicated genotypes fed NCD (n=6/group). (J) *Proglucagon* mRNA levels in ileum of 14- to 16-week-old male mice of the indicated genotypes fed NCD. (n=6/group). (K) The plasma GLP-1 levels in 14- to 16-week-old male mice of the indicated genotypes fed NCD (n=6/group). (L) Representative images for Piezo1 RNA-FISH and GLP-1 immunofluorescent staining in the ileum of 14-week-old male mice of indicated genotypes fed NCD (n=6/group). (M) Percentage of Piezo1-positive GLP-1 cells in total GLP-1 cells in the ileal mucosa of 14-week-old male mice of indicated genotypes fed NCD (n=6/group). (N) A schematic diagram depicting the potential mechanisms linking the CaMKKβ/CaMKIV-mTOR signaling pathway and GLP-1 production. (O) Representative western blots are shown for indicated antibodies in the ileal mucosa (n = 6/group). Data are represented as mean ± SEM. Significance was determined by Student’s t test for comparison between two groups, and by one-way ANOVA for comparison among three groups or more, *p < 0.05, **p < 0.01, ***p < 0.001.

Under normal chow diet, *IntL-Piezo1^-/-^* mice exhibited increased body weight (Figure 1G) and greater glycemic excursions compared to control groups (*Piezo1^loxp/loxp^*, *Vil-Flp*, *FGC* and *IntL-cre*) (Figure 1H and I), while the food and water intake were not changed (Figure 1-figure supplement 2A and B). The morphology of islet (Figure 1-figure supplement 3A) and ileum (Figure 1-figure supplement 4A) were not affected. Ileal mucosal Proglucagon expression and plasma GLP-1 level were significantly lower in *IntL-Piezo1^-/-^* mice than that in all littermate controls such as *Piezo1^loxp/loxp^*, *Vil-Flp*, *FGC* and *IntL-Cre* mice (Figure 1J and K), while no significant alteration was observed in the expression of pancreatic Piezo1 and Proglucagon (Figure 1-figure supplement 3B-D). According to in situ hybridization of Piezo1 and GLP-1, the expression of Piezo1 disappeared in GLP-1 positive cells, suggesting successful knockout of Piezo1 in L cells in *IntL-Piezo1^-/-^*mice (Figure 1L and M). Also depicted in Figure 1-figure supplement 5, Piezo1 is expressed in GLP-1-positive cells of the duodenum, jejunum, ileum, and colon in control mice, but not in *IntL-Piezo1^-/-^* mice. However, Piezo1 remains expressed in intestinal ghrelin positive cells and pancreatic glucagon positive cells of *IntL-Piezo1^-/-^* mice (Figure 1-figure supplement 6). Moreover, while GLP-1 levels were reduced in L cells of *IntL-Piezo1^-/-^* mice, levels of PYY, another hormone secreted by L cells, were unaffected (Figure 1-figure supplement 7A-D). Additionally, ileal mucosal cholecystokinin (CCK), a hormone secreted by I cells with metabolic effects similar to GLP-1, was also unchanged in *IntL-Piezo1^-/-^* mice (Figure 1-figure supplement 7E). Previous study showed that Piezo1 affected intestinal tight junctions and epithelial integrity (Jiang et al., 2021). To access whether loss of Piezo1 in L cells affect epithelial integrity of the intestine, we examined the expression of tight junction proteins, including ZO-1 and Occludin. As shown in Figure 1-figure supplement 8, the expression of ZO-1 and Occludin remained unchanged in *IntL-Piezo1^-/-^*mice when compared to littermate controls.

Piezo1 is a non-selective cationic channel that allows passage of Ca^2+^ and Na^+^. CaMKKβ is the main calcium/calmodulin dependent protein kinase kinase involved in the regulation of metabolic homeostasis (Marcelo et al., 2016). It is activated by binding calcium-calmodulin (Ca^2+^/CaM), resulting in downstream activation of kinases CaMKIV. The activation of CaMKIV modulate the gene expression of nutrient- and hormone-related proteins (Ban et al., 2000; Chen et al., 2011; Takemoto-Kimura et al., 2017). Previous studies have reported that Ca^2+^ and mTOR signaling regulate the production of GLP-1 (Tolhurst et al., 2011; Xu et al., 2015; Yu & Jin, 2010). Drawing from these findings, we hypothesized that Piezo1 might regulate GLP-1 synthesis through the CaMKKβ/CaMKIV-mTOR signaling pathway (Figure 1N). As shown in Figure 1O, abrogated GLP-1 production was associated with decreased CaMKKβ/CaMKIV-mTOR signaling in the ileal mucosa of *IntL-Piezo1^-/-^*mice (Figure 1O).

### Derangements of glucose metabolism and GLP-1 production were induced by HFD in *IntL-Piezo1^-/-^* mice, which was mitigated by Exendin-4

We next assessed the effect of L cell-specific *Piezo1* gene deletion on GLP-1 and glucose tolerance in diet-induced diabetic mice. *IntL-Piezo1^-/-^* and control mice were exposed to HFD for 10 weeks. Compared to the controls, higher body weight (Figure 2A), greater glucose excursions (Figure 2B) were observed in *IntL-Piezo1^-/-^* mice exposed to HFD. Ileal mucosal Proglucagon expression levels were lower in *IntL-Piezo1^-/-^* than control mice (Figure 2C-F). Impaired CaMKKβ/CaMKIV-mTORC1 signaling pathway in ileal mucosa as evidenced by a decrease in CaMKKβ, reduced phosphorylation levels of CaMKIV, mTOR, S6K, and S6 was also observed in *IntL-Piezo1^-/-^*mice (Figure 2F). No significant alteration in morphology, Piezo1 or Proglucagon levels were observed in the pancreas of *IntL-Piezo1^-/-^*mice (Figure 2-figure supplement 3E-H). Together these data demonstrate that *IntL-Piezo1^-/-^* mice with prolonged HFD feeding exhibit impaired glucose metabolism phenotype and reduced GLP-1. Injection of GLP-1 analog Exendin-4 (Ex-4) decreased the body weight (Figure 2G) and improved both glucose tolerance (Figure 2H) and insulin resistance (Figure 2I) in control and *IntL-Piezo1^-/-^* mice, while endogenous synthesis of GLP-1 was not changed by Ex-4 injection in *IntL-Piezo1^-/-^* mice (Figure 2J and K). These data suggested that decreased GLP-1 synthesis and secretion contribute to impaired glucose metabolism in *IntL-Piezo1^-/-^* mice.

**Figure 2:**
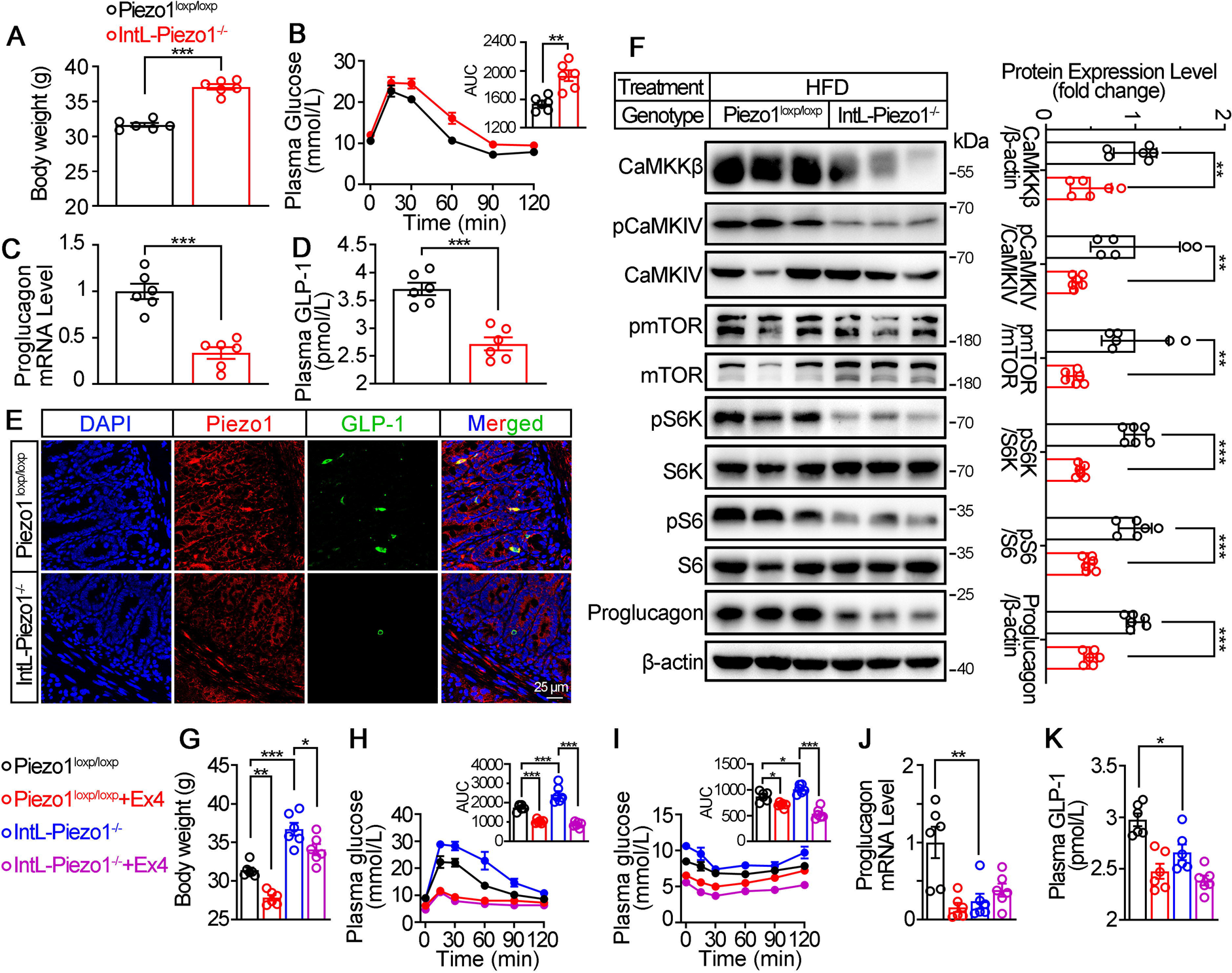
Validation and phenotype of *IntL-Piezo1^-/-^* mice fed with high-fat diet. (A) Body weight of 14- to 16-week-old male *Piezo1^loxp/loxp^* and *IntL-Piezo1^-/-^* mice fed with HFD for 10 weeks (n=6/group). (B) IPGTT and associated area under the curve (AUC) values of 14- to 16-week-old male *Piezo1^loxp/loxp^* and *IntL-Piezo1^-/-^* mice fed with HFD (n=6/group). (C) *Proglucagon* mRNA levels in the ileal mucosa of 14- to 16-week-old male *Piezo1^loxp/loxp^* and *IntL-Piezo1^-/-^* mice fed with HFD (n=6/group). (D) The plasma GLP-1 level in 14- to 16-week-old male *Piezo1^loxp/loxp^* and *IntL-Piezo1^-/-^* mice fed with HFD (n=6/group). (E) Double immunofluorescent staining of Piezo1, and GLP-1 in the ilea of 14- to 16-week-old male *Piezo1^loxp/loxp^* and *IntL-Piezo1^-/-^* mice fed HFD (n=6/group). (F) Representative western blots are shown for indicated antibodies in the ileal mucosa (n=6/group). (G) Body weight after 7 consecutive days infusion of saline or Ex-4 (100ug/kg body weight) in 14- to 16-week-old male *Piezo1^loxp/loxp^*and *IntL-Piezo1^-/-^* mice fed with HFD (n=6/group). (**H**-**I**) IPGTT (**H**) and ITT (**I**) and associated area under the curve (AUC) values after consecutive infusion of saline or Ex-4. (J) *Proglucagon* mRNA levels in the ileal mucosa (n=6/group) after consecutive infusion of saline or Ex-4. (K) The plasma GLP-1 level after consecutive infusion of saline or Ex-4 (n=6/group). Data are represented as mean ± SEM. Significance was determined by Student’s t test for comparison between two groups, and by one-way ANOVA for comparison among three groups or more, *p < 0.05, **p < 0.01, ***p < 0.001.

### The pharmacological and mechanical activation of ileal Piezo1 stimulates GLP-1 synthesis

We next examined whether activation of Piezo1 could rescued the impaired glucose metabolism in diet-induced diabetic mice. Injection of Piezo1 activator Yoda1 after 10 weeks of high-fat diet, led to reduced body weight and improved the impaired glucose metabolism significantly in diabetic mice, while Piezo1 antagonist GsMTx4 reversed the weight loss and glucose-lowering effect of Yoda1 (Figure 3A and B). Yoda1 remarkably induced an increase in GLP-1 synthesis and secretion (Figure 3C and D), as well as an increment of CaMKKβ/CaMKIV-mTORC1 signaling in ileal mucosa (Figure 3E), while GsMTx4 abolished the effect of Yoda1 (Figure 3C-E). However, weight loss, improved plasma glucose and increased GLP-1 production induced by Yoda1 were not observed in *IntL-Piezo1^-/-^* mice (Figure 3F-J).

**Figure 3:**
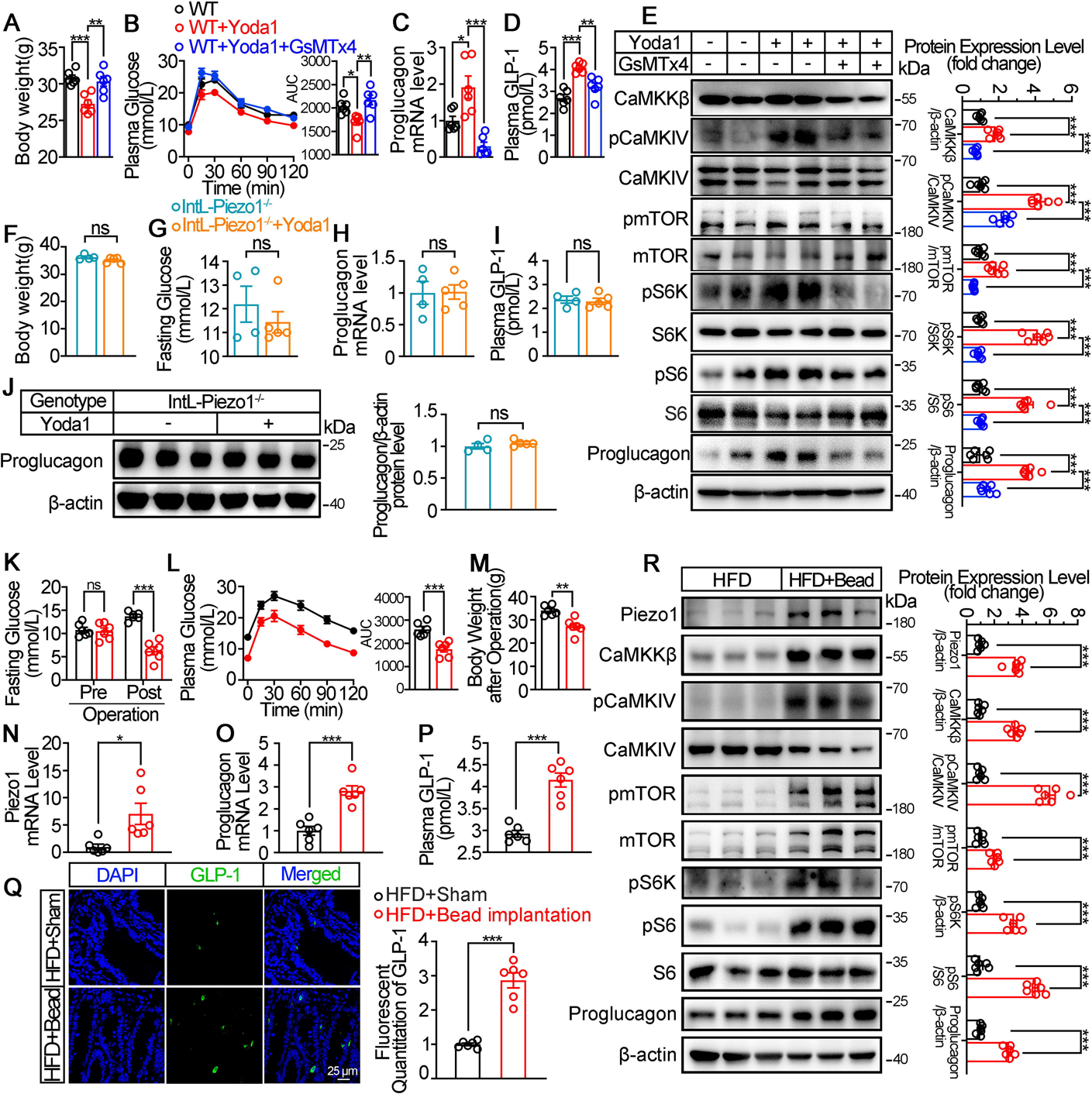
Chemical and mechanical interventions of Piezo1 regulates GLP-1 synthesis in mice. (**A**-**E**) 14- to 16-week-old male C57BL/6J mice fed with HFD for 10 weeks were infused with vehicle, Yoda1 (2 μg per mouse) or GsMTx4 (250 μg/kg) by i.p. for 7 consecutive days. (n=6/group). (A) Body weight after consecutive drug infusion. (B) IPGTT and associated area under the curve (AUC) values. (C) *Proglucagon* mRNA levels in the ileal mucosa. (D) Plasma GLP-1. (E) Representative western blots are shown for indicated antibodies in the ileal mucosa (**F**-**J**) 14- to 16-week-old male *IntL-Piezo1^-/-^* mice fed with HFD for 10 weeks were infused with vehicle, Yoda1 (2 μg per mouse) by i.p. for 7 consecutive days. (n=4 or 5/group) (F) Body weight after 7 consecutive days’ drug infusion. (G) Fasting blood glucose levels. (H) Ileal mucosal *Proglucagon* mRNA levels. (I) Plasma GLP-1 levels. (J) Ileal mucosal Proglucagon protein levels. (**K-R**) 14- to 16-week-old male C57BL/6J mice fed with HFD were subjected to sham operation, or intestinal bead implantation (n=6/group). (K) Fasting blood glucose levels. (L) IPGTT and associated area under the curve (AUC) values. (M) Body weight. (**N** and **O**) *Piezo1* (**N**) and *Proglucagon* (**O**) mRNA levels in the ileal mucosa. (P) The plasma GLP-1 levels. (Q) Immunofluorescence staining of GLP-1 in ileum and quantification of GLP-1 positive cells. (R) Representative western blots images and densitometry quantification for indicated antibodies in the ileal mucosa. Data are represented as mean ± SEM. Significance was determined by Student’s t test for comparison between two groups, and by one-way ANOVA for comparison among three groups or more, *p < 0.05, **p < 0.01, ***p < 0.001.

The intestine receives mechanical stimulation from the chyme, which may activate Piezo1 in the intestine epithelium, including L cells. To mimic the mechanical pressing and stretching induced by intestinal contents, a small silicon bead was implanted into the high-fat diet-induced diabetic mouse ileum. To exclude the possibility of bowel obstruction and abdominal pain caused by bead implantation, we measured the fecal mass and gastrointestinal transit time, and accessed abdominal mechanical sensitivity in both sham and bead-implanted mice. As shown in Figure3-figure supplement 9A-B, there was no significant difference in fecal mass and gastrointestinal transit time between the sham-operated mice and those implanted with beads. The results of abdominal mechanical sensitivity indicated that no difference in abdominal pain threshold was observed between sham and bead implanted mice (Figure3-figure supplement 9C). Intestinal bead implantation improved the impaired glucose metabolism in diabetic mice (Figure 3K and L). Body weight loss, activated ileal mucosal CaMKKβ/CaMKIV-mTOR signaling, increased mRNA and protein levels of ileal mucosal Piezo1 and Proglucagon, as well as the circulating levels of GLP-1 were observed in diabetic mice after operation (Figure 3M-R). The above data suggest that mechanical stimuli induced by intestinal bead implantation activates ileal Piezo1 in diabetic mice, stimulating GLP-1 production via CaMKKβ/CaMKIV-mTOR signaling axis, thus improving glucose homeostasis.

### Piezo1 regulates GLP-1 synthesis and secretion in primary cultured mouse L cells and isolated mouse ileum

To obtain primary L cells, we isolated cell from the ileum of *IntL-Cre-mT/mG* mice, in which tdTomato expression switched to EGFP expression in L cells as shown in Figure 1E. EGFP positive cells (mouse L cells) were then sorted from isolated single cells (Figure 4A). Immunofluorescence showed that the sorted EGFP+ cells were Piezo1 positive (Figure 4B).

**Figure 4:**
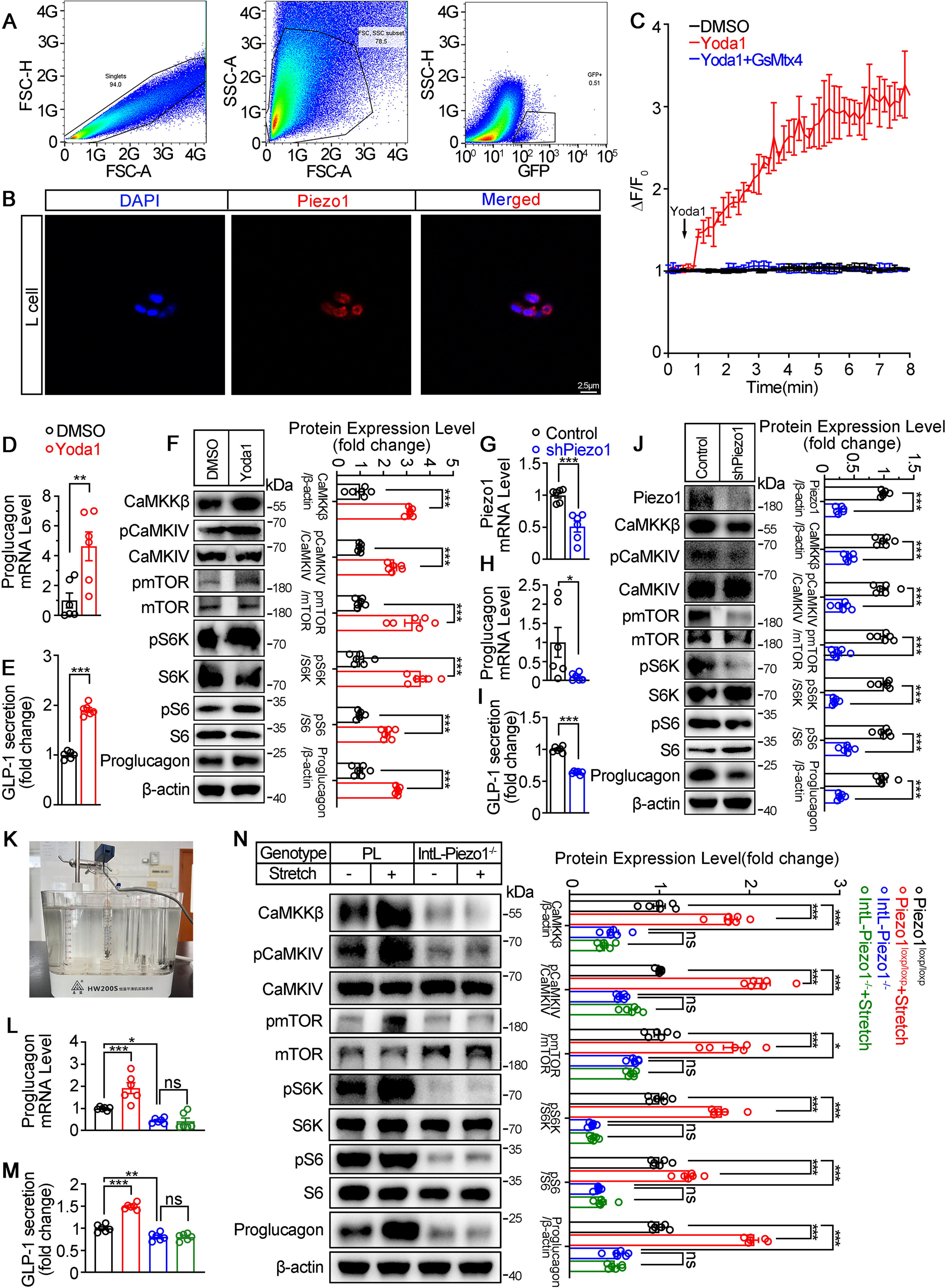
Piezo1 regulates GLP-1 synthesis and secretion in primary cultured mouse L cells and isolated mouse ileum. (A) Isolation of mouse L cells (GFP positive) from ileal tissue by FACS. The gating in flowcytometry for sorting of GFP positive cells. (B) Immunofluorescent staining of Piezo1 in sorted GFP positive L cells. (C) Intracellular Ca^2+^ imaging by fluo-4-AM calcium probe. The change of fluorescent intensity (ΔF/F0) was plotted against time (**D**-**F**) L cells were treated with vehicle or Yoda1 (5μM) for 24 hours. (D) *Proglucagon* mRNA expression. (E) GLP-1 concentrations in the culture medium. (F) Western blot images and densitometry quantification for the indicated antibodies. (**G**-**J**) Knockdown of Piezo1 in L cells by shRNA for 48 hours. (G) *Piezo1* mRNA expression. (H) *Proglucagon* mRNA expression. (I) GLP-1 levels in the culture medium. (J) Western blot images and densitometry quantification for the indicated antibodies. (**K**-**N**) Ileal tissues from *Piezo1^loxp/loxp^* and *IntL-Piezo1^-/-^* mice were subjected to tension force (n=6/group). (K) A representative photograph showing the traction of isolated ileum. (L) *Proglucagon* mRNA levels. (M) GLP-1 concentrations in the medium. (N) Western blot images and densitometry quantification for the indicated antibodies. Data are represented as mean ± SEM and are representative of six biological replicates. Significance was determined by Student’s t test for comparison between two groups, and by one-way ANOVA for comparison among three groups or more, *p < 0.05, **p < 0.01, ***p < 0.001.

Yoda1 at the dose of 5μM triggered an increase in intracellular Ca^2+^ level in primary cultured mouse L cells, which was blocked by pre-incubation of cells with GsMTx4 (0.1 μM) for 15 min (Figure 4C). Yoda1 also stimulated Proglucagon expression and GLP-1 secretion, as well as CaMKKβ/CaMKIV-mTOR signaling pathway in primary cultured mouse L cells (Figure 4D-F). In contrast, knockdown of Piezo1 by shRNA led to significant decrease in Proglucagon expression and GLP-1 secretion, as well as inhibition of CaMKKβ/CaMKIV/mTOR signaling pathway (Figure 4G-J).

Given the ability of Piezo1 in sensing mechanical force, tension of 1.5g was applied to the isolated mouse ileum bathed in Tyrode’s solution for two hours. Tension stimulated Proglucagon expression, GLP-1 secretion and activated CaMKKβ/CaMKIV-mTOR signaling pathway in the ileum of control mice, but not in *IntL-Piezo1^-/-^* mice (Figure 4K-N), suggesting the involvement of Piezo1 of the L cells in mediating the force-induced GLP-1 production and CaMKKβ/CaMKIV-mTOR signaling.

### Pharmacological, mechanical and genetic activation of Piezo1 stimulates GLP-1 synthesis and secretion in STC-1 cells

To further validate the role of Piezo1 in regulating GLP-1, we examined the effect of manipulating Piezo1 on GLP-1 production in an intestinal neuroendocrine cell line STC-1. Pharmacological activation of Piezo1 by Yoda1 triggered an inward current in STC-1 cell recorded by whole cell patch-clamp, which could be inhibited by pre-incubation of GsMTx4 (Figure 5A). Yoda1 also triggered an increase in intracellular Ca^2+^ level in STC-1 cells. Pre-incubation of cells with GsMTx4 (0.1 μM) for 15 min inhibited [Ca^2+^]_i_ increase (Figure 5B and C). Yoda1 induced a concentration-dependent activation of CaMKKβ/CaMKIV-mTOR pathway and GLP-1 synthesis and secretion (Figure 5D-F). GsMTx4 blocked the effect of Yoda1 on STC-1 cells in both GLP-1 and CaMKKβ/CaMKIV-mTOR activation (Figure 5G-I).

**Figure 5:**
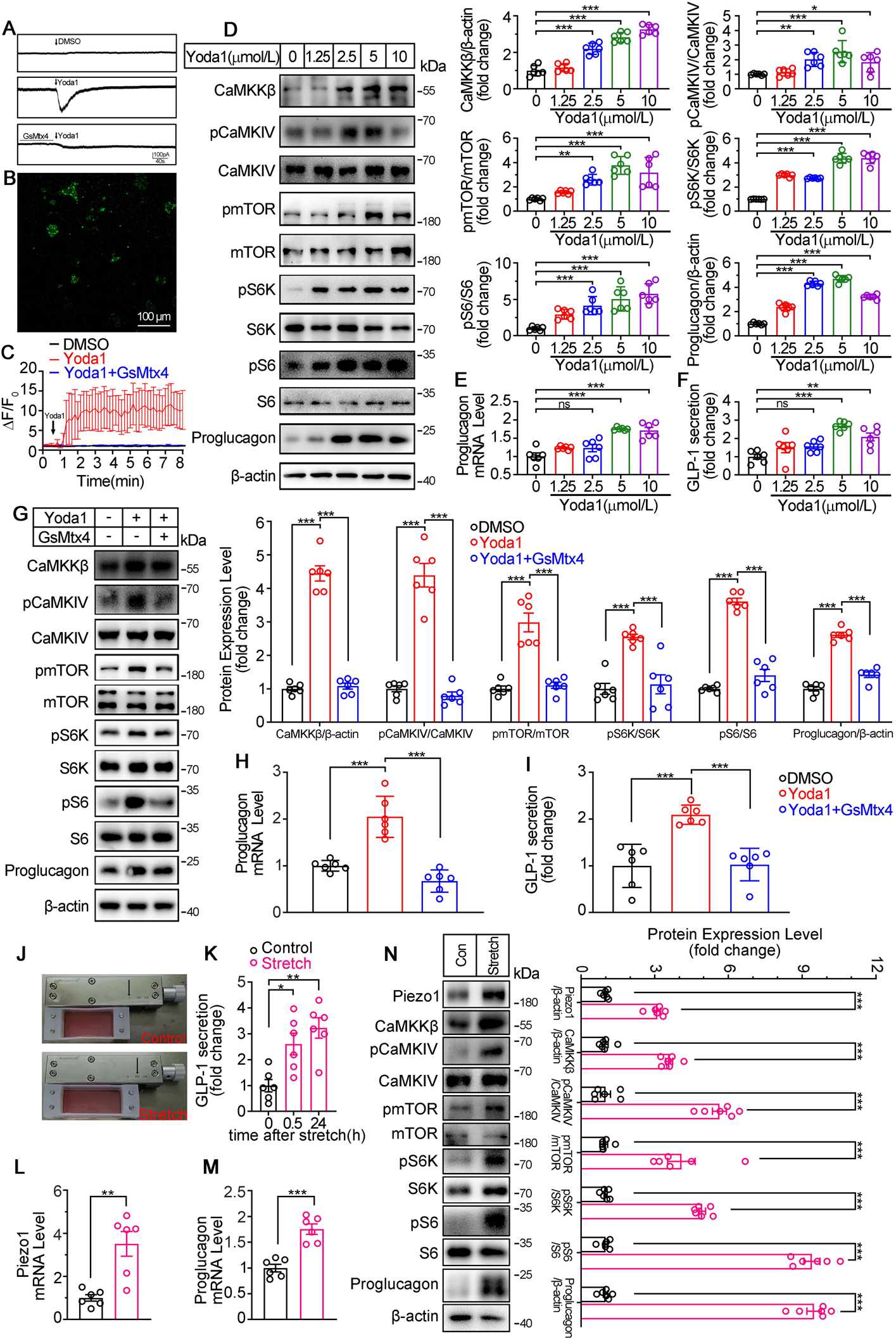
Modulation of GLP-1 synthesis and secretion by pharmacological and mechanical activation of Piezo1 in STC-1 cells. (A) Whole-cell currents induced by Yoda1 (5μM) were recorded from STC-1 cells or STC-1 cells pretreated with GsMTx4 for 30 min (**B** and **C**) Intracellular calcium imaging in STC-1 cells. (**B**) STC-1 cells were loaded with fluo-4 AM for 1 h. The representative time-lapse image showing the intracellular Ca^2+^ signals. (**C**) The change of fluorescent intensity (ΔF/F0) was plotted against time. (**D-F**) STC-1 cells were treated with various concentrations of Yoda1 for 24 h. (**D**) Whole-cell extracts underwent western blot with indicated antibodies. (**E**) *Proglucagon* mRNA levels. (**F**) GLP-1 concentrations in the culture medium. (**G-I**) STC-1 cells were treated with Yoda1 (5 μM) in the presence or absence of GsMTx4 (0.1 μM) for 24 h. (**G**) Whole-cell extracts underwent western blot with indicated antibodies. (**H**) *Proglucagon* mRNA levels. (**I**) GLP-1 concentrations in the culture medium (**J-N**) STC-1 were subjected to mechanical stretch. (**J**) STC-1 cells were cultured in elastic chambers and the chambers were subjected to mechanical stretch by 120% extension of their original length. (**K**) The medium GLP-1 concentrations were detected at indicated time. (**L**) *Piezo1* mRNA levels. (**M**) *Proglucagon* mRNA levels. (**N**) Whole-cell extracts underwent western blot with indicated antibodies. Data are represented as mean ± SEM and are representative of six biological replicates. Significance was determined by Student’s t test for comparison between two groups, and by one-way ANOVA for comparison among three groups or more, *p < 0.05, **p < 0.01, ***p < 0.001.

To mimic the activation of Piezo1 by mechanical stretching in vivo, STC-1 cells grown on elastic chambers were subjected to mechanical stretch to 120% of their original length. Mechanical stretch upregulated Piezo1 and Proglucagon expression, promoted GLP-1 secretion (Figure 5J-N), and activated CaMKKβ/CaMKIV- mTOR signaling pathways (Figure 5N).

Consistent to the pharmacological and mechanical activation of Piezo1, over-expression of Piezo1 in STC-1 cells resulted in a significant increase in GLP-1 production, as well as activation of the CaMKKβ/CaMKIV-mTOR signaling pathway (Figure 6A-D). Conversely, knockdown of Piezo1 by shRNA led to a significant decrease in GLP-1 production and inhibition of CaMKKβ/CaMKIV-mTOR signaling pathways (Figure 6E-H).

**Figure 6:**
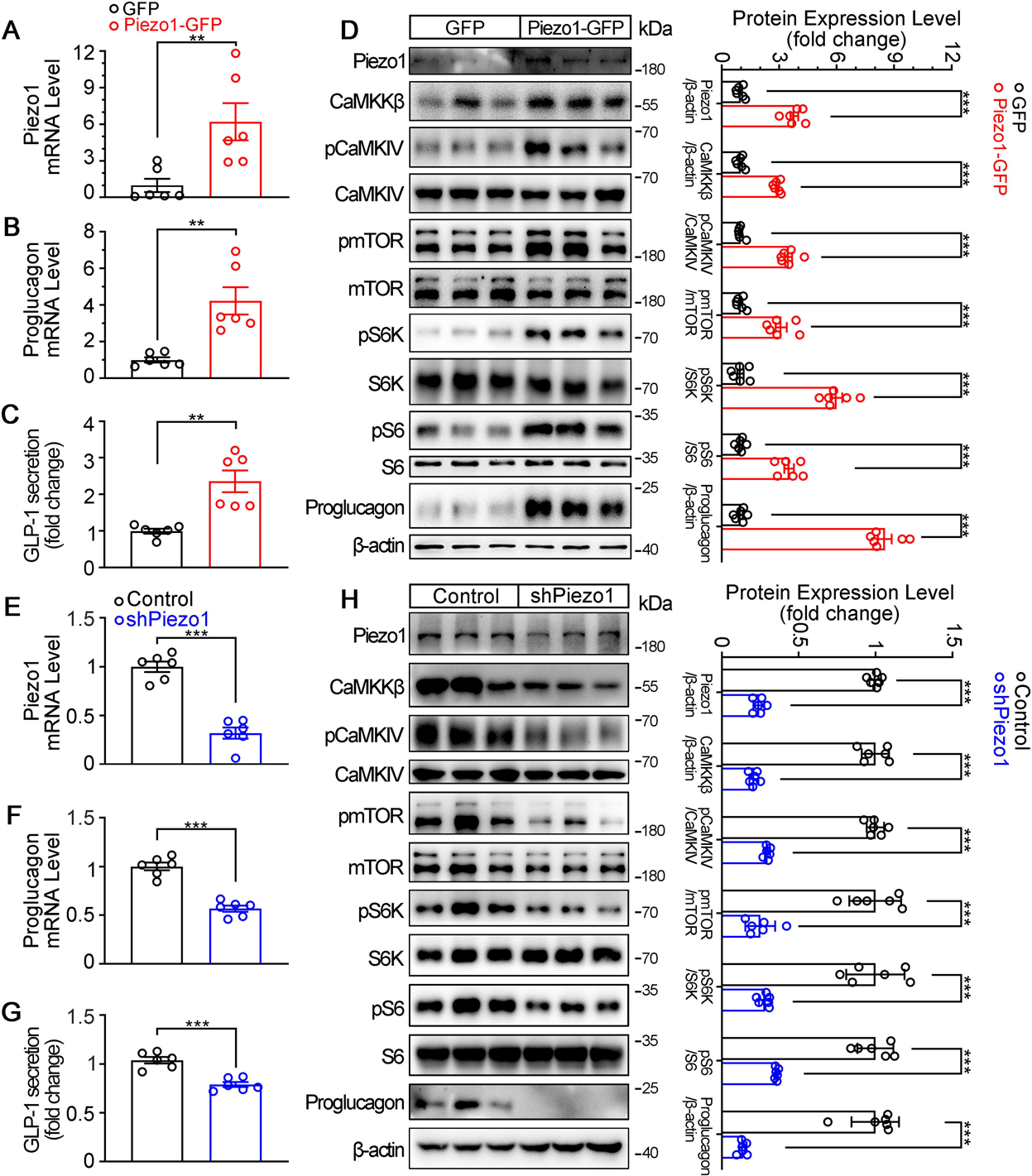
Genetic interference of Piezo1 regulates GLP-1 production in STC-1 cells. (**A-D**) STC-1 cells were transfected with mouse control or *Piezo1* expression plasmids for 48h. *Piezo1* (**A**) and *Proglucagon* (**B**) mRNA levels in STC-1 cells. (**C**) GLP-1 concentrations in culture medium. (**D**) Whole-cell extracts underwent western blot with indicated antibodies. (**E**-**H**) Stable knockdown of Piezo1 in STC-1 cells. *Piezo1* (**E**) and *Proglucagon* (**F**) mRNA levels in STC-1 cells. (**G**) GLP-1 concentrations in culture medium. (**H**) Whole-cell extracts underwent western blot with indicated antibodies. Data are represented as mean ± SEM Data are represented as mean ± SEM and are representative of six biological replicates. Significance was determined by Student’s t test, *p < 0.05, **p < 0.01, ***p < 0.001.

### Piezo1 regulates GLP-1 production through CaMKK**β**/CaMKIV and mTOR in STC-1 cells

Next, we examined whether CaMKKβ/CaMKIV and mTOR signaling mediates the effects of Piezo1 on GLP-1 production. Overexpression of CaMKKβ or CaMKIV increased CaMKKβ/CaMKIV and mTOR signaling activity, resulting in increased synthesis and secretion of GLP-1 (Figure 7A-C). In contrast, the CaMKKβ inhibitor STO-609, downregulated CaMKKβ/CaMKIV and mTOR signaling, as well as GLP-1 synthesis and secretion (Figure 7D-F). Inhibition of mTORC1 activity by rapamycin suppressed GLP-1 production induced by Yoda1, which was associated with inhibition of mTOR signaling (Figure 7G-I).

**Figure 7:**
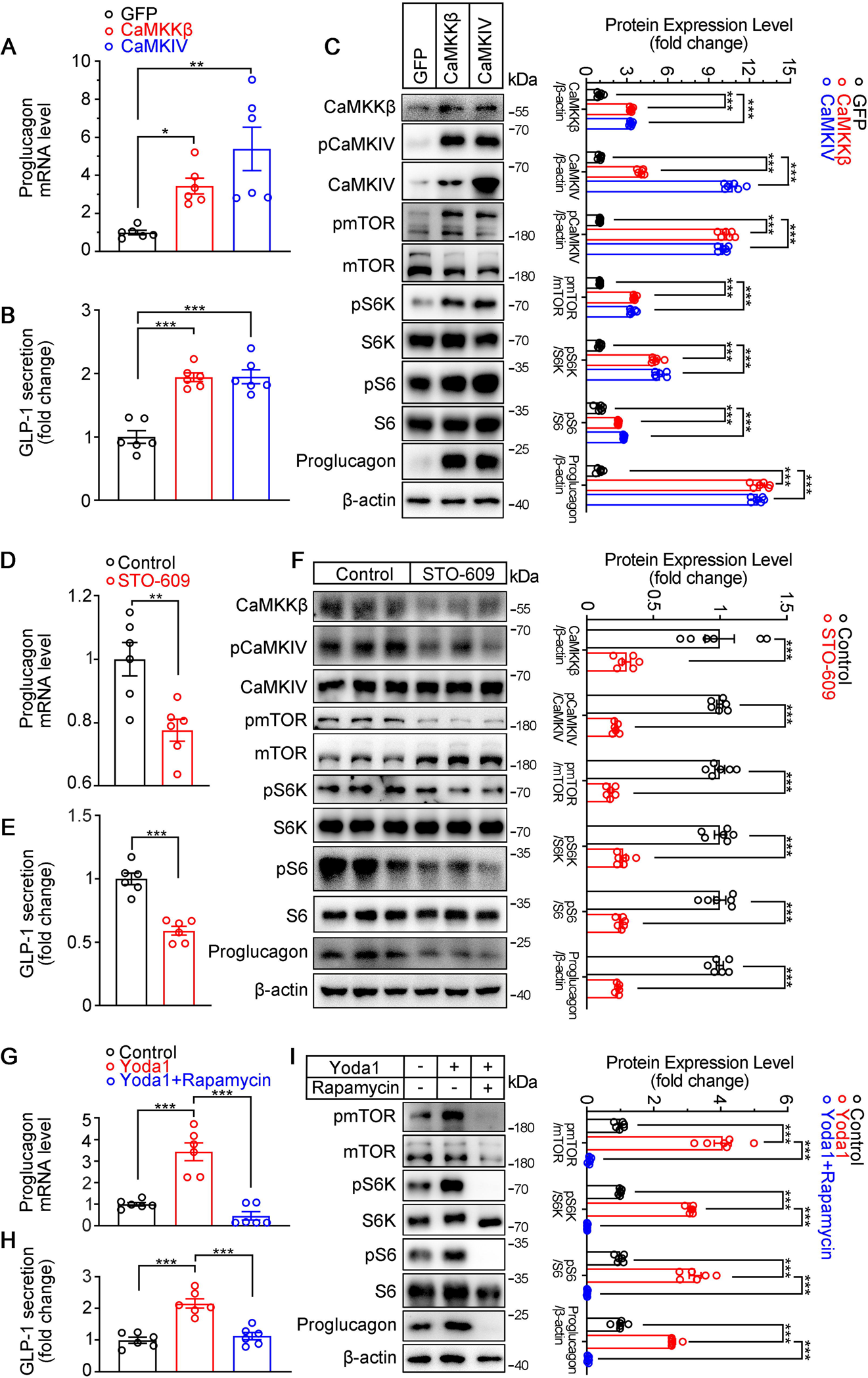
Modulation of GLP-1 production by CaMKKβ/CaMKIV and mTOR signaling activity in STC-1 cells. (**A-C**) STC-1 cells were transfected with GFP, CaMKKβ or CaMKIV plasmids for 48h. (**A**) *Proglucagon* mRNA levels in STC-1 cells. (**B**) GLP-1 concentrations in culture medium. (**C**) Whole-cell extracts underwent western blot with indicated antibodies. (**D-F**) STC-1 cells were treated with CaMKKβ inhibitor STO-609 (10 μmol/L) for 24 h. (**D**) *Proglucagon* mRNA levels in STC-1 cells. (**E**) GLP-1 concentrations in culture medium. (**F**) Whole-cell extracts underwent western blot with indicated antibodies. (**G-I**) STC-1 cells were pretreated with Rapamycin (50 nmol/L) for 1 h, then treated with Yoda1 (5 μmol/L) for 24 h. (**G**) *Proglucagon* mRNA levels in STC-1 cells. (**H**) GLP-1 concentrations in the culture medium. (**I**) Whole-cell extracts underwent western blot with indicated antibodies. Data are represented as mean ± SEM Data are represented as mean ± SEM and are representative of six biological replicates. Significance was determined by Student’s t test for comparison between two groups, and by one-way ANOVA for comparison among three groups or more, *p < 0.05, **p < 0.01, ***p < 0.001.

## Discussion

It has been known for decades that GLP-1 secretion from the intestinal L cells is stimulated by meal intake and is essential for postprandial glycemic control (Drucker, 2006; Song et al., 2019). However, the mechanism underlying the regulation of GLP-1 production is not completely understood. One of the problems that impeded the investigation of regulation mechanism of GLP-1 is the lack of an L cell-specific genetically engineered animal model. Here, we generated an L cell-specific Cre mouse line for the first time by combination of the Flp-Frt and Cre-LoxP systems, which allows genetic manipulation specifically in the L cells and thus creates a useful tool to investigate molecular mechanisms in L cells.

Previous studies have shown that L cells are able to sense nutrients in the intestinal lumen such as glucose and other carbohydrates, lipids and amino acids, which induce GLP-1 secretion through different mechanisms, including membrane depolarization-associated exocytosis, Ca^2+^/Calmodulin (Tolhurst et al., 2011), cAMP (Yu & Jin, 2010), mTORC1 (Xu et al., 2015) and AMPK (Jiang et al., 2016) signaling pathways. However, it is innegligible that as open type endocrine cells, L cells not only receive the chemical stimulations from the nutrients, but also mechanical stimulation when the chyme passing through the intestine, including stretching, pressure and shear force (Sensoy, 2021). While the food needs to be digested and nutrients absorbed before L-cells can detect the nutritive signals, mechanical stimulation may be more direct and faster. Here, we showed the expression of the mechano-sensitive ion channel Piezo1 in L cells of human and mouse intestinal sections, mouse primary L cell culture and an intestinal neuroendocrine cell line STC-1, suggesting the mechano-sensing ability of L cells and potential regulation of GLP-1 in response to mechanical stimulation. Indeed, our study showed that the mechano-regulation of GLP-1 secretion did exist as demonstrated by the increased GLP-1 secretion by intestinal bead implantation, intestinal tissue stretching or STC-1 cell stretching. Moreover, mice with selective loss of Piezo1 expression in intestinal L cell (*IntL-Piezo1^-/-^*) exhibited reduced circulating levels of GLP-1, increased body weight and impaired glucose homeostasis, while pharmacological activation of Piezo1 on mice, primary L cells and STC-1 cells did the reverse. More importantly, *IntL-Piezo1^-/-^* mice was unable to response to the tension-induced GLP-1 production. These further suggested a Piezo1-mediated mechanical sensing mechanism in L cells that regulates GLP-1 production and glucose metabolism by sensing the stimulation of intestinal luminal contents. Therefore, our study provides a mechano-regulation mechanism in addition to the existing known nutritive regulation for GLP-1 production. However, the relationship between mechano-regulation and nutritive regulation remains to be explored.

Interestingly, this intestinal Piezo1-mediated mechanical sensing mechanism may severely impaired in diabetic patients and rodents. We observed a decreased Piezo1 expression in the ileal mucosa of diet-induced diabetic mice accompanied by reduced GLP-1 production. When challenged with high-fat, *IntL-Piezo1^-/-^* mice exhibited more severe symptoms of diabetes which was mitigated by Ex-4. These findings suggest that the impairment of Piezo1-mediated mechanical sensing function in the intestine is an important mechanism for the pathogenesis of T2DM. It is noteworthy that RYGB, a commonly performed weight-loss and hypoglycemic surgery (Cummings et al., 2004), significantly increased Piezo1 expression in L cells of obese diabetic patients. Yoda1 treatment or intestinal bead implantation enhanced GLP-1 production and improved glucose metabolism in the diet-induced diabetic mouse model, suggesting that restore the mechano-sensing or enhance the function of Piezo1 either pharmacologically or mechanically, may be a new strategy to improve the secretion of GLP-1, thus alleviate T2DM. However, in our study, the Piezo1-mediated regulation of GLP-1 production is only demonstrated in transgenic mice, mouse primary L cells and an intestinal neuroendocrine cell line derived from mouse. Whether Piezo1 plays the same role in human L cells awaits to be investigated. A number of studies have generated L cells culture from human intestinal organoid culture or human intestinal stem cell monolayer culture by manipulating the growth factors in the media (Goldspink et al., 2020; Petersen et al., 2014; Villegas-Novoa et al., 2022). It is worthy to validate our finding in human L cells in order to prove its translational potential in T2DM treatment.

The intragastric balloon is a current noninvasive clinical weight loss measure that involves placing a space-occupying balloon in the stomach to reduce food intake and generate satiety signals, thus maintaining satiety. Investigations illustrated that intragastric balloon alter the secretion of hormones such as cholecystokinin and pancreatic polypeptide, delay the emptying of food in the stomach and reduce the appetite (Mathus-Vliegen & de Groot, 2013). Intragastric balloon provides a feasible weight loss intervention for obese people (Kim et al., 2016). In this study, a new intestinal implantation surgery of beads was adopted, which may offer a novel approach for weight loss and glucose control by activating the intestinal Piezo1-GLP-1 signaling pathway in the future.

Mechanistically, our study suggest that Piezo1 regulates GLP-1 production through a CaMKKβ/CaMKIV-mTOR signaling pathway. CaMKKβ/CaMKIV has been reported to mediate the Ca^2+^ signaling in many metabolic processes, including liver gluconeogenesis and de novo lipogenesis, adipogenesis, insulin sensitivity and β cell proliferation (Anderson et al., 2012; Lin et al., 2011; Liu et al., 2012; J. Liu et al., 2022). mTOR plays a central role in nutrient and energy sensing and regulates cellular metabolism and growth in response to different nutrient and energy status (Howell & Manning, 2011). Here we showed that mTOR can also response to mechanical stimuli through a mechano-sensitive Ca^2+^ channel mediated CaMKKβ/CaMKIV activation. Although we did not demonstrate direct phosphorylation of mTOR or S6K by CaMKIV in L cells, previous study reported that CaMKKβ could serve as a scaffold to assemble CaMKIV with key components of the mTOR/S6K pathway and promote liver cancer cell growth (Lin et al., 2015), which lend support to the CaMKKβ/CaMKIV-mTOR signaling in our study. Recently, Knutson et. al. found that ryanodine and IP3-triggered calcium release from intracellular calcium store could amplified the initial Peizo2 - Ca^2+^ signal triggered by mechanical stimulation, and was required for the mechanotransduction in the serotonin release from enterochromaffin cells (Knutson et al., 2023). In our study, we also observed long-lasting intracellular Ca^2+^ increase triggered by Yoda1 in primary L cells and STC-1 cells, which also suggested an involvement of intracellular Ca^2+^ store in the Ca^2+^ relay. Beside Ca^2+^, cyclic AMP (cAMP) is another signaling molecule that active *Gcg* gene expression and GLP-1 production (Drucker et al., 1994; Jin, 2008; Simpson et al., 2007). cAMP was found to play a critical role in nutrients-induced GLP-1 secretion, including glucose (Ong et al., 2009), lipids (Hodge et al., 2016), and amino acids (Tolhurst et al., 2011). Previous study reported that Ca^2+^ can activate soluble adenylyl cyclase (sAC) to increase intracellular cAMP (Jaiswal & Conti, 2003). Whether sAC-cAMP can be activated by Piezo1-mediated Ca^2+^ influx and whether it is an alternative signaling pathway that mediates the Piezo1-regulated GLP-1 production remain to be explored.

In summary, our study reveals a previously undiscovered Piezo1-mediated mechano-sensing property of intestinal L cell, which plays an essential role in the regulation of GLP-1 production and glucose metabolism. This finding also suggests a new mechano-regulation in enteroendocrine cells, in addition to chemical and neuronal regulation, which may shed new light on the strategy for metabolic diseases such as diabetes and obesity.

## Additional information

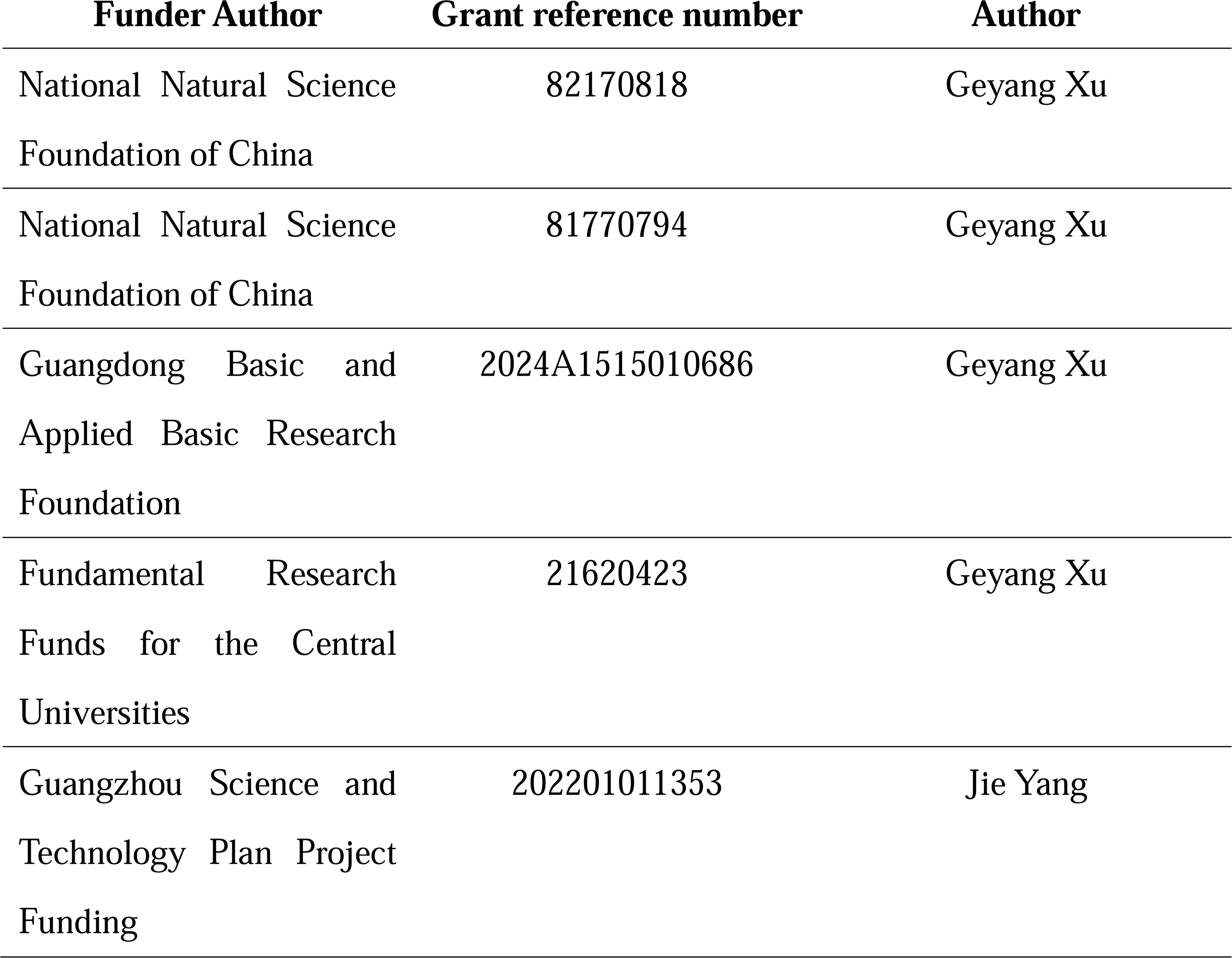

## Author contribution

Yanling Huang, Data curation, Software, Formal analysis, Validation, Investigation, Methodology, Writing - original draft;

Haocong Mo, Data curation, Software, Formal analysis, Validation, Investigation, Methodology, Writing - original draft;

Jie Yang, Data curation, Formal analysis, Validation, Investigation, Methodology, Funding acquisition;

Luyang Gao, Data curation, Formal analysis, Validation, Investigation, Methodology, Writing - original draft;

Tian Tao, Data curation, Investigation, Methodology; Qing Shu, Formal analysis, Validation, Investigation; Wenying Guo, Formal analysis, Validation, Investigation; Yawen Zhao, Formal analysis, Validation, Investigation; Jingya Lyu, Resources, Formal analysis, Validation; Qimeng Wang, Formal analysis, Validation;

Jinghui Guo, Resources, Investigation; Hening Zhai, Resources, Investigation;

Linyan Zhu, Resources, Investigation, Methodology;

Hui Chen, Conceptualization, Resources, Formal analysis, Supervision, Investigation, Software, Visualization, Methodology, Writing - review and editing;

Geyang Xu, Conceptualization, Resources, Formal analysis, Supervision, Funding acquisition, Investigation, Visualization, Methodology, Writing - review and editing, Project administration.

## Ethical Statement

Animals used in this study were handled in accordance with the Guide for the Care and Use of Laboratory Animals published by the National Institutes of Health (NIH Publications No. 8023, revised 1978). All animal protocols were approved by the Animal Care and Use Committee of Jinan University.

Human subject research was approved by the Institutional Review Board of Jinan University.

## Data availability

All of the data supporting the findings of this study are included in the article and supplementary information.

## Abbreviations

CaMKKβ: Ca^2+^/calmodulin-dependent protein kinase kinase β
CaMKIV: Ca^2+^/calmodulin-dependent protein kinases IV
CCK: Cholecystokinin
EECs: Enteroendocrine cells
mTOR: Mechanistic target of rapamycin
PYY: Peptide tyrosine tyrosine
RYGB: Roux-en-Y gastric bypass
S6K: p70 S6 Kinase
S6: S6 Ribosomal Protein
T2DM: Type 2 Diabetes Mellitus

