## Supplementary information figures for "Mechano-regulation of GLP-1 production by Piezo1 in intestinal L cells"

### Supplementary Methods and Materials

#### Detection of abdominal mechanical sensitivity

The mice were familiarized with a metal mesh floor covered with plastic boxes for 2 hours each day for 2 days prior to testing. Their abdominal fur was shaved 1 day before the experiments. The abdominal area was then stimulated using calibrated von Frey filaments (VFFs) that applied varying forces: subthreshold mechanical stimuli (indicative of allodynia, 0.07 g force) and painful stimuli (indicative of hyperalgesia, 0.16 and 1 g force). Each filament was applied 10 times for 5-8 seconds with 10-second intervals between applications. To prevent learning or sensitization, the same area was not stimulated more than once consecutively. The data were presented as the number of withdrawal responses out of 10 applications, with 0 indicating no withdrawal and 10 indicating the maximum withdrawal. A withdrawal response was defined as (1) the mouse withdrawing its abdomen from the VFFs, (2) subsequent licking of the abdominal area, or (3) withdrawal of the entire body. All tests were conducted in a blinded manner.

#### Immunofluorescence

Paraffin-embedded tissue sections were dewaxed and rehydrated, followed by boiling in 0.01 mol/L citrate buffer (pH 6.0) for 10 minutes. Subsequently, the sections were blocked with 5% goat serum for 1 hour and then incubated overnight at 4°C with the following primary antibodies: Piezo1 (1:400), Glucagon (1:200), Ghrelin (1:100), GLP-1(1:500), PYY (1:100), ZO-1 (1:200), or Occludin (1:200). After the primary antibody incubation, the sections were exposed to secondary antibodies for 2 hours at room temperature. Finally, fluorescence micrographs were captured using a confocal laser scanning microscope (Leica M205FA, Germany).

### Gastrointestinal Transit Time

The whole-gut transit time test was conducted as previously described (Qin et al., 2017). The duration between the oral administration of charcoal and the appearance of the first stained fecal pellet was recorded as the total gastrointestinal transit time.

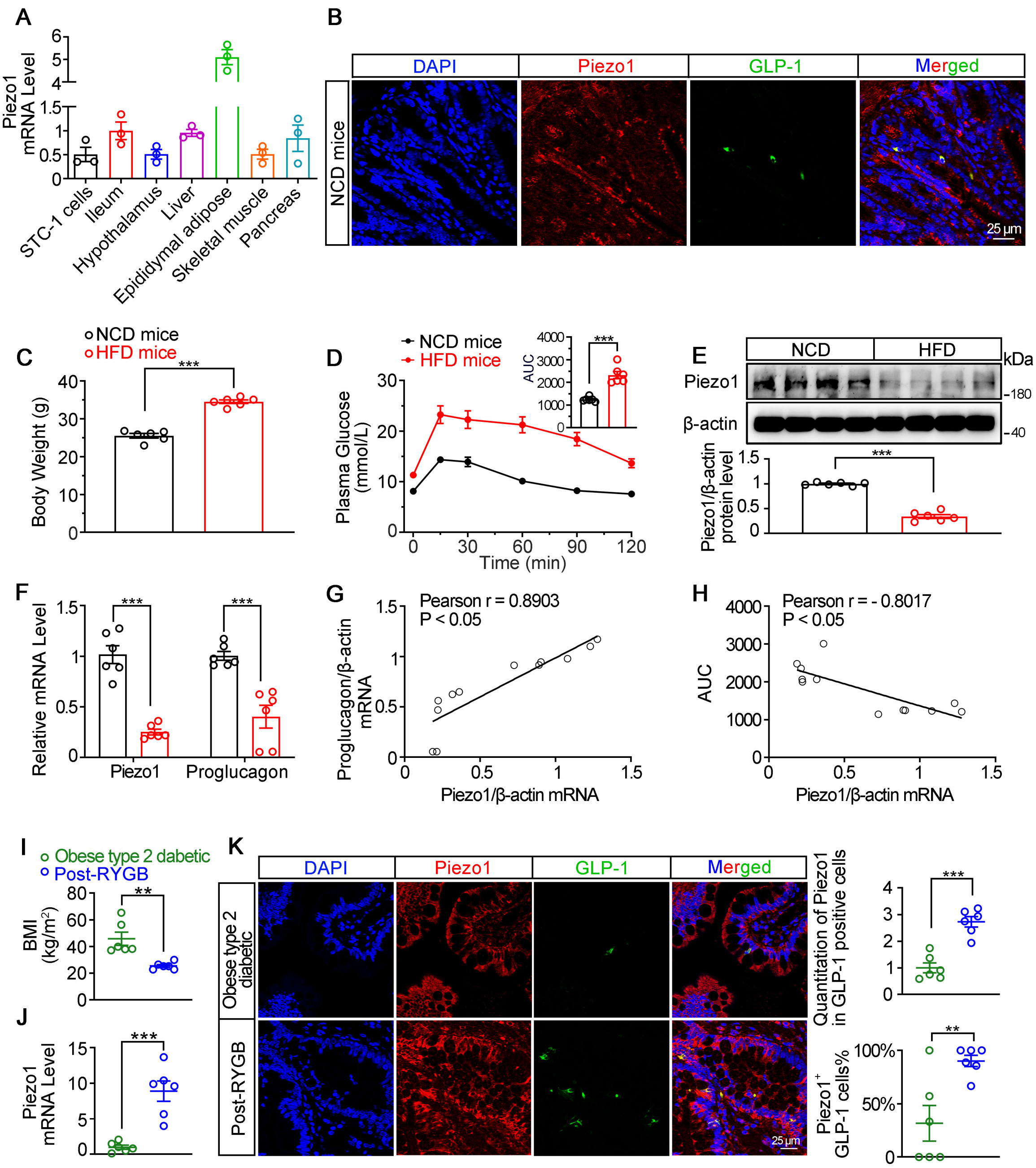


**Figure supplement 1: Assessment of Piezo1 and GLP-1 in mouse and human ilea.**

(**A**) *Piezo1* mRNA levels in STC-1 cells and various tissues of 14-week-old male C57BL/6J mice fed normal chow diet (NCD).

(**B**) Double immunofluorescent staining of Piezo1 (red) and GLP-1 (green) in the NCD mouse ileum.

(**C**) Body weight of 14-week-old male C56BL/6J mice were fed with either normal chow diet (NCD) or high fat diet (HFD) (n=6/group).

(**D**) IPGTT and associated area under the curve (AUC) values of 14-week-old male C56BL/6J mice fed NCD or HFD (n=6/group).

(**E**) Representative western blots are shown for Piezo1 and β-actin protein levels in the ileal mucosa of 14-week-old male C56BL/6J mice fed NCD or HFD (n=6/group).

(**F**) *Piezo1* and *Proglucagon* mRNA levels in the ileal mucosa of 14-week-old male C56BL/6J mice fed NCD or HFD detected by qPCR (n=6/group).

(**G**) Pearson's correlation analysis of the correlation between ileal mucosal *Piezo1* and *Proglucagon* mRNA levels in 14-week-old male C56BL/6J mice fed a NCD or HFD.

(**H**) Pearson's correlation analysis of the correlation between area under the curve (AUC) for glucose excursion and ileal mucosal *Piezo1* mRNA level in 14-week-old male C56BL/6J mice fed NCD or HFD.

(**I**) Body mass index (BMI) of post-RYGB subjects and obese type 2 diabetics (n = 6/group).

(**J**) *Piezo1* mRNA levels in the ileal mucosa of post-RYGB subjects and obese type 2 diabetics by qPCR (n = 6/group).

(**K**) Double immunofluorescent staining of Piezo1 and GLP-1 in the ileum of post-RYGB patients and obese type 2 diabetic patients. (n = 6/group).

Data are represented as mean ± SEM. Significance was determined by Student's t test for comparison between two groups, and by one-way ANOVA for comparison among three groups or more. *$p < 0.05$, **$p < 0.01$, ***$p < 0.001$.

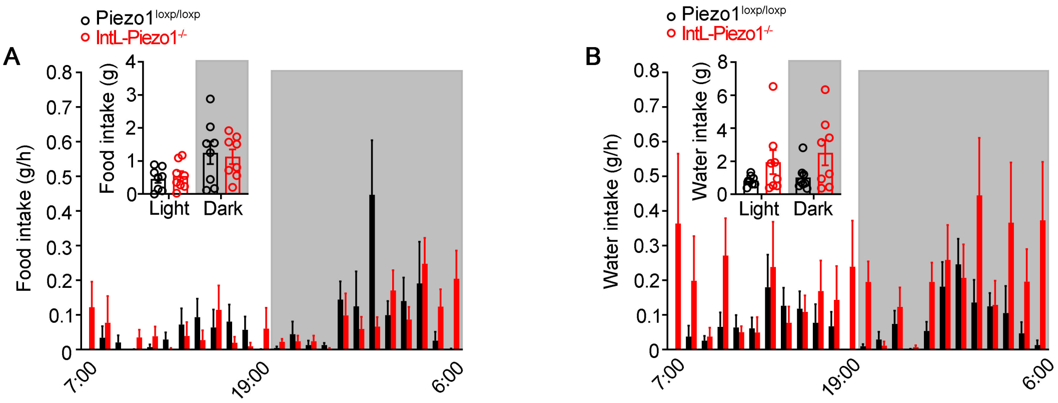


**Figure supplement 2: Food intake and water intake of *IntL-Piezo1$^{-/-}$* mice.**
(**A**) Food intake and (**B**) water intake of 12- to 14-week-old male *Piezo1$^{loxp/loxp}$* and *IntL-Piezo1$^{-/-}$* mice fed with normal chow diet (n=8/group).
Data are represented as mean ± SEM. Significance was determined by Student's t test. *p < 0.05, **p < 0.01, ***p < 0.001.

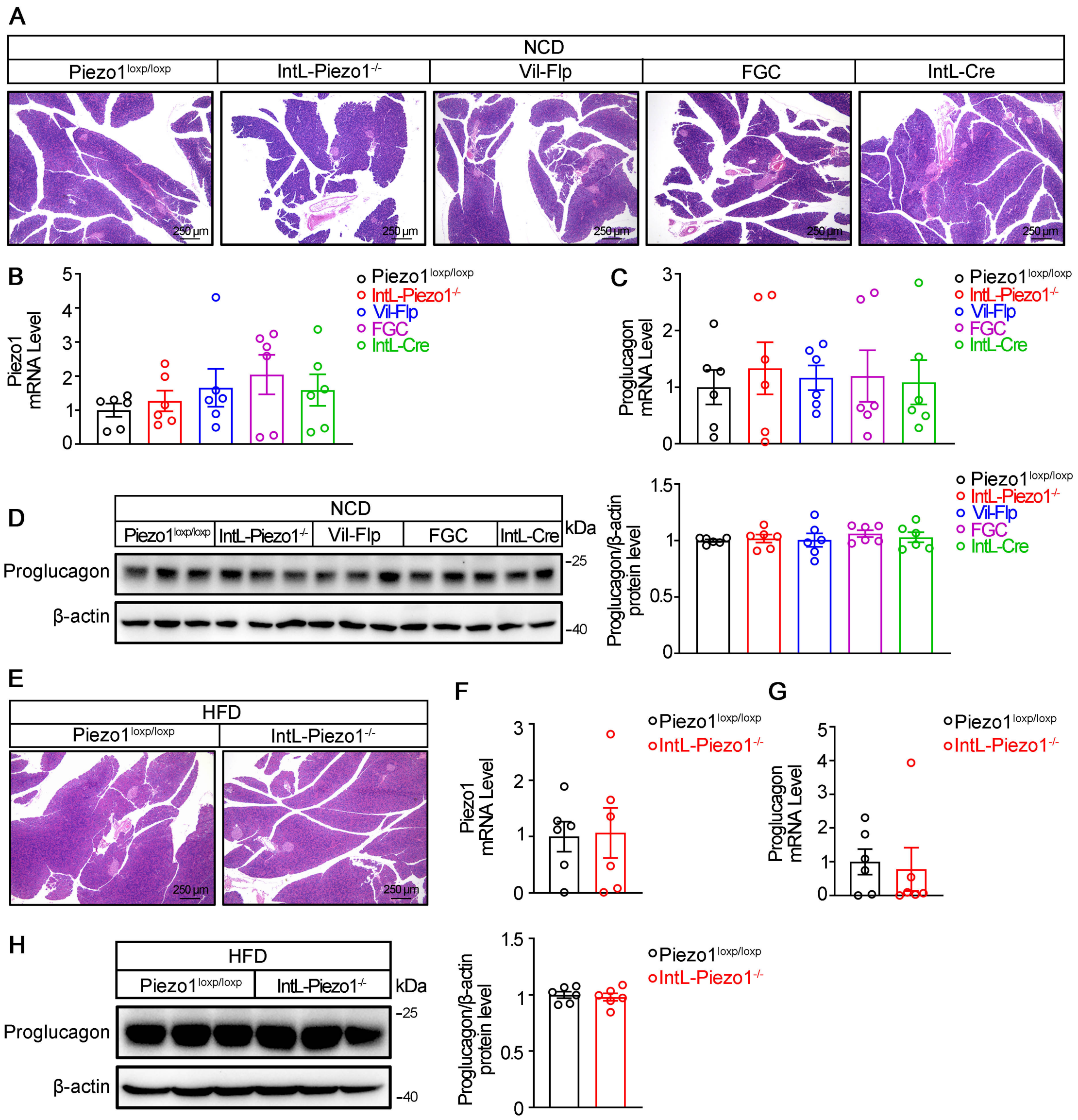


**Figure supplement 3: *IntL-Piezo1$^{-/-}$* mice preserve normal pancreatic morphology and proglucagon expression.**

(**A**) HE staining of pancreatic sections from 14- to 16-week-old male mice of the indicated genotypes fed with NCD.

(**B** and **C**) *Piezo1* (**B**) and *Proglucagon* (**C**) mRNA levels in pancreas of 14- to 16-week-old male mice of the indicated genotypes fed with NCD. (n=6/group).

(**D**) Western blot analysis of Proglucagon protein levels in pancreas of 14- to 16-week-old male mice of the indicated genotypes fed NCD (n=6/group).

(**E**) HE staining of pancreatic sections from 14- to 16-week-old male *Piezo1$^{loxp/loxp}$* and *IntL- Piezo1$^{-/-}$* mice fed HFD.

(**F** and **G**) *Piezo1* (**F**) and *Proglucagon* (**G**) mRNA levels in pancreas of 14- to 16-week-old male *Piezo1$^{loxp/loxp}$* and *IntL- Piezo1$^{-/-}$* mice fed HFD (n=6/group).

(**H**) Western blot analysis of Proglucagon protein levels in pancreas of 14- to 16-week-old male *Piezo1$^{loxp/loxp}$* and *IntL- Piezo1$^{-/-}$* mice fed HFD (n=6/group).

Data are represented as mean ± SEM. Significance was determined by Student's t test for comparison between two groups, and by one-way ANOVA for comparison among three groups or more, *$p < 0.05$, **$p < 0.01$, ***$p < 0.001$.

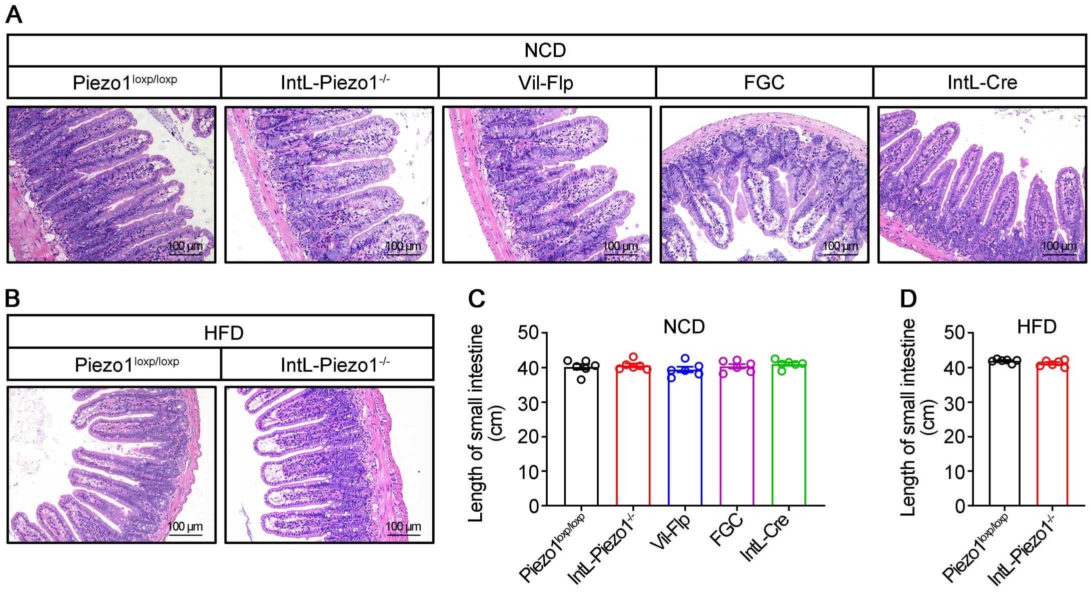


**Figure supplement 4: Intestinal morphology of *IntL-Piezo1*$^{-/-}$ mice.**

(**A** and **B**) HE staining of ileal sections from 14- to 16-week-old male mice of the indicated genotypes fed with NCD (**A**) or HFD (**B**).

(**C** and **D**) The length of small intestine from male mice of the indicated genotypes fed NCD (**C**) or HFD (**D**) (n=6/group).

Data are represented as mean ± SEM. Significance was determined by Student's t test for comparison between two groups, and by one-way ANOVA for comparison among three groups or more.

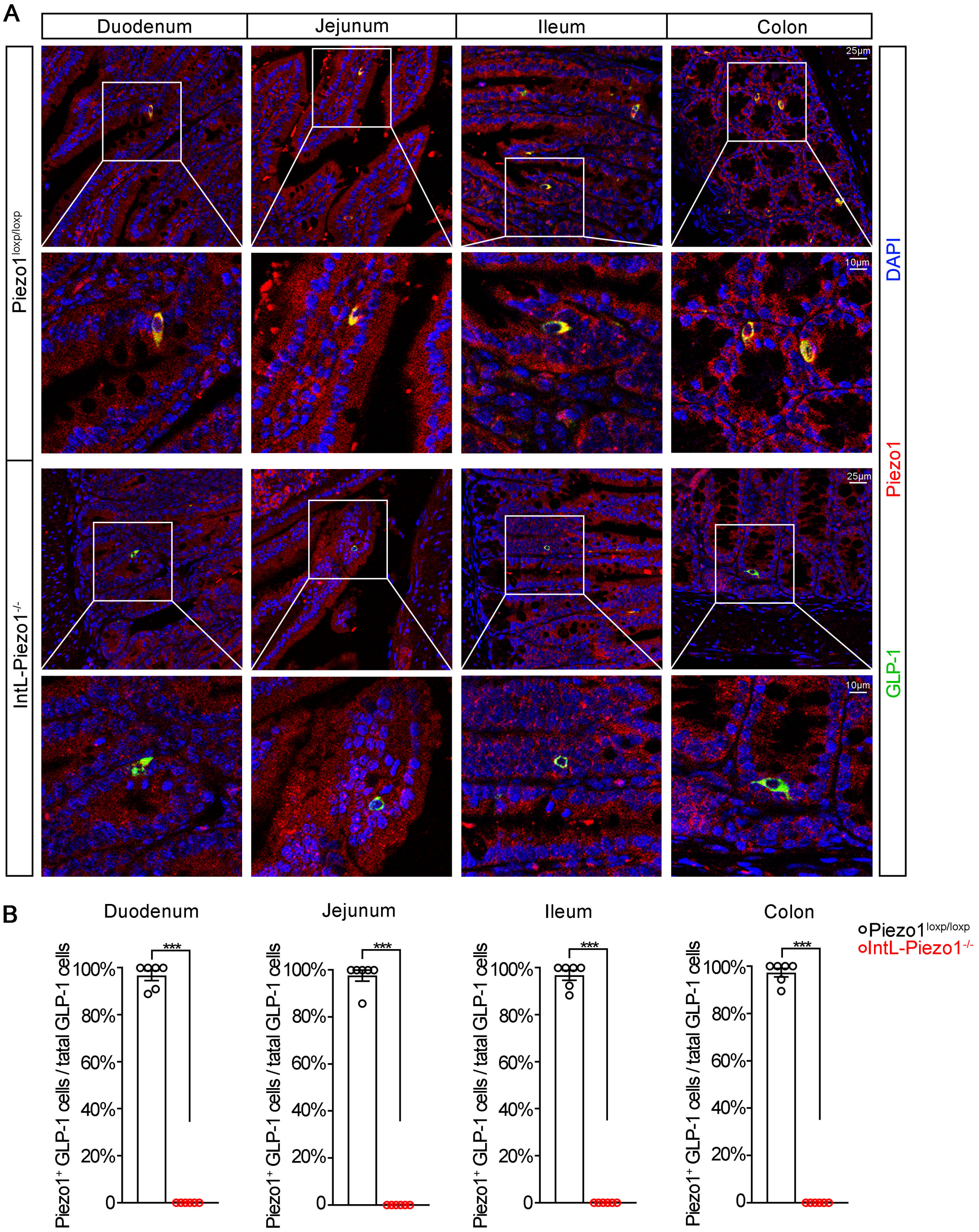


**Figure supplement 5: Double immunostaining of Piezo1 and GLP-1 in the intestines of *IntL-Piezo1$^{-/-}$* mice.**

(**A**) Representative images for Piezo1 and GLP-1 immunofluorescent staining from different regions of the intestine of 14-week-old male mice of the indicated genotypes fed NCD (n=6/group).

(**B**) Percentage of Piezo1-positive GLP-1 cells in total GLP-1 cells in the different regions of intestinal mucosa of 14-week-old male mice of the indicated genotypes fed NCD (n=6/group).

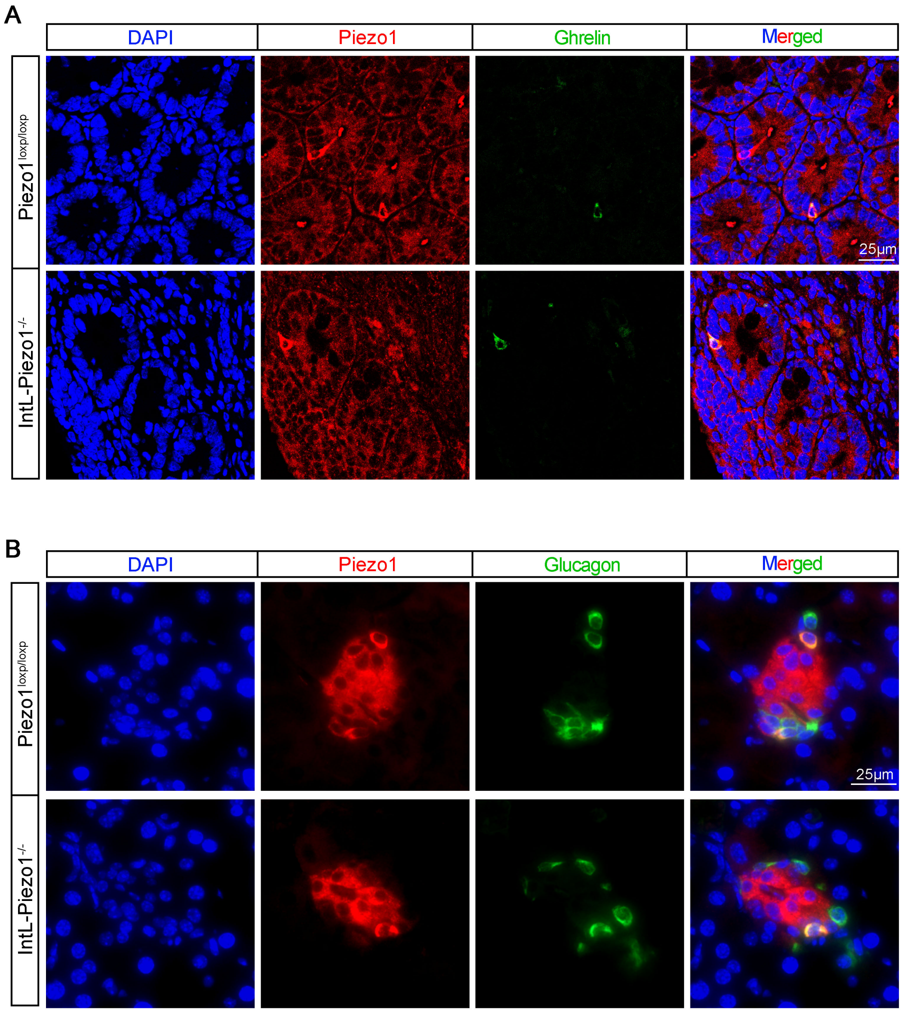


**Figure supplement 6: Expression of Piezo1 in intestinal ghrelin cells and pancreatic α cells .**

(**A**) Representative images for Piezo1 and Ghrelin immunofluorescent staining in the ileum of 14-week-old male mice of the indicated genotypes fed NCD (n=6/group).

(**B**) Representative images for Piezo1 and Glucagon immunofluorescent staining in the pancreas of 14-week-old male mice of the indicated genotypes fed NCD (n=6/group).

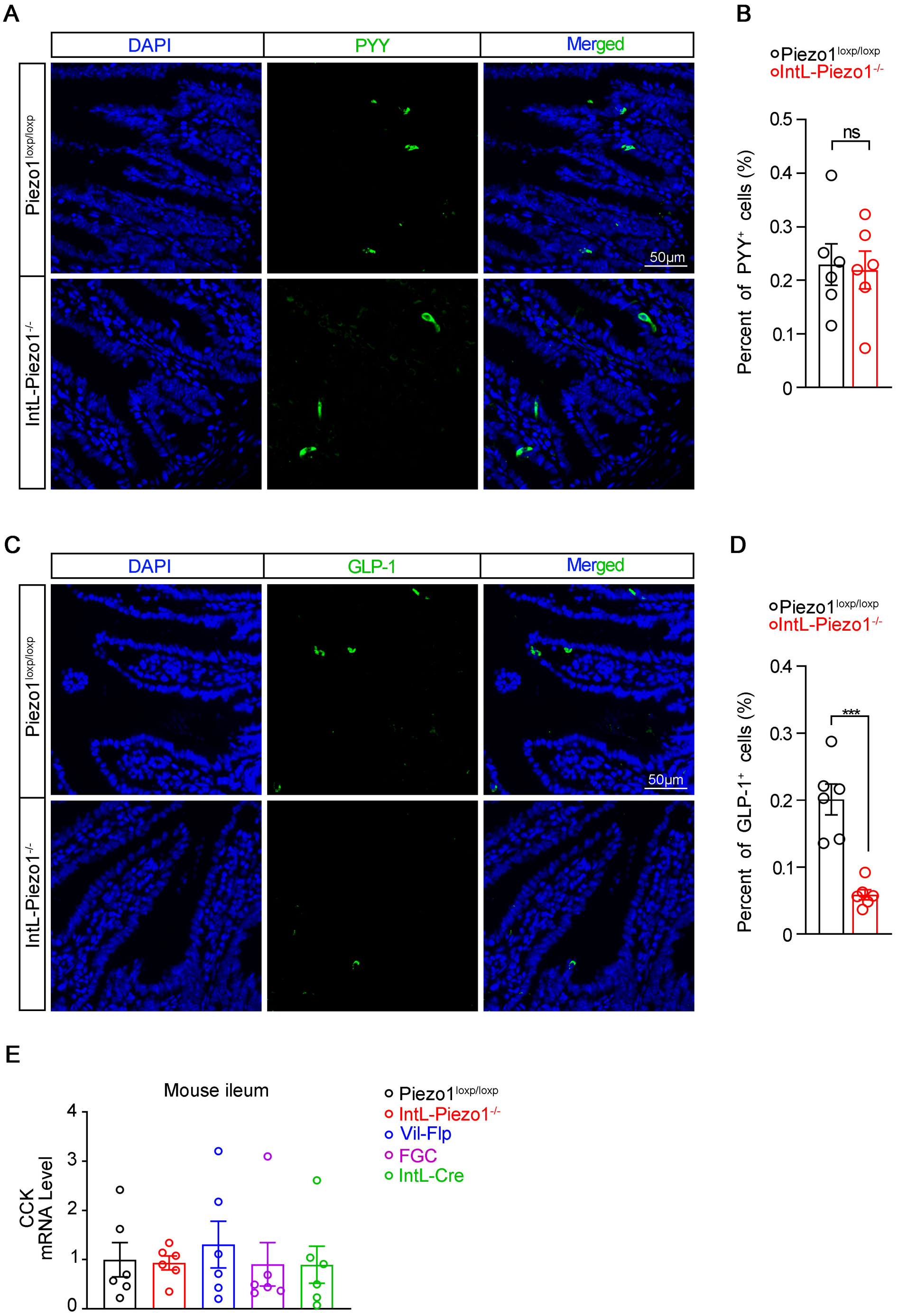


**Figure supplement 7: Assessment of L cell hormones and CCK in the ileum of *IntL-Piezo1*$^{-/-}$ mice.**

(**A**) Representative images for Peptide YY (PYY) immunofluorescent staining in the ileum of 14-week-old male mice of the indicated genotypes fed NCD (n=6/group).

(**B**) Percentage of PYY positive cells in ileal mucosal cells (n=6/group).

(**C**) Representative images for GLP-1 immunofluorescent staining in the ileum of 14-week-old male mice of the indicated genotypes fed NCD (n=6/group).

(**D**) Percentage of GLP-1 positive cells in ileal mucosal cells (n=6/group).

**(E)** Ileal mucosal CCK mRNA levels of 14- to 16-week-old male mice of the indicated genotypes fed with NCD (n=6/group).

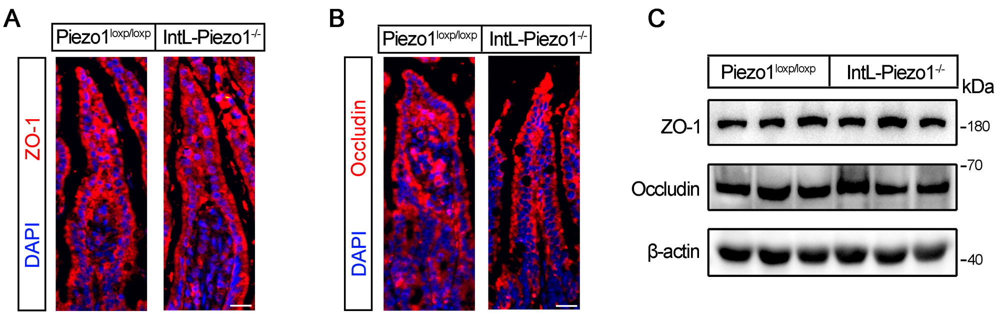


**Figure supplement 8: Effect of L cell-specific Piezo1 deletion on intestinal barrier function and tight junction proteins.**

(**A**) Representative images for ZO-1 immunofluorescent staining in the ileum of 14-week-old male mice of the indicated genotypes fed NCD (n=6/group).

(**B**) Representative images for Occludin immunofluorescent staining in the ileum of 14-week-old male mice of the indicated genotypes fed NCD (n=6/group).

(**C**) Representative western blots are shown for ZO-1, Occludin and β-actin protein levels in the ileal mucosa of 14-week-old male C56BL/6J mice fed NCD or HFD (n=6/group).

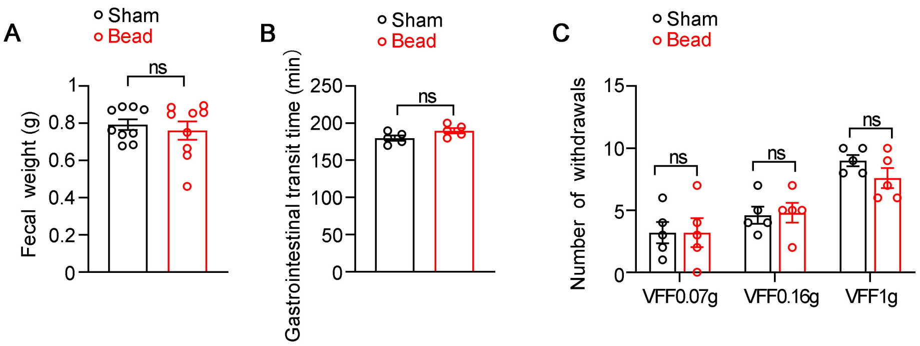


**Figure supplement 9: Effect of intestinal bead implantation on fecal weight, gastrointestinal transit time and abdominal pain in C57BL/6J mice.**

(**A**) Fecal weight of sham and bead implanted mice fed HFD. (n=9/group).

(**B**) Gastrointestinal transit time of sham and bead-implanted mice fed HFD. (n=5/group).

(**C**) Assessment of abdominal mechanical sensitivity. Mechanical sensitivity of the abdomen was assessed using calibrated von Frey filaments (0.07 g, 0.16 g, and 1 g) in sham and bead-implanted mice (n=5 per group).
