## Supplementary information tables for "Mechano-regulation of GLP-1 production by Piezo1 in intestinal L cells"

**Supplementary information, Table S1. Sequences of the genotyping primers,** **Related to experimental model and subject details**

| **Primers** | **Sequence (5’→3’)** | **Primer type** |
| --- | --- | --- |
| P1 | GACCTTTGCCCTCTGGTCTC | Forward |
| P2 | GAGTGACGGTGCCAGAGAAA | Reverse |
| P3 | GACTCCAGCTGCCTTCTCTG | Forward |
| P4 | CGGTGATCTCCCAGATGCTC | Reverse |
| P5 | CCCTAACTCAGTCTCCAGCA | Forward |
| P6 | CGGTTACCAGGTGGTCATGT | Reverse |
| P7 | CCCTAACTCAGTCTCCAGCA | Forward |
| P8 | CTGCAAAGGGTCGCTACAGA | Reverse |
| P9 | AATGGCTCTCCTCAAGCGTAT | Forward |
| P10 | ACAGGAGGTAGTCCCTCACAT | Reverse |
| P11 | TGTCGGGGAAATCATCGTCC | Reverse |

**Supplementary information, Table S2. Sequences of primers used in RT-PCR experiments,** **Related to method details**

|  | **Upstream primer (5’-3’)** | **Downstream primer (5’-3’)** | **Accession number(s)** |
| --- | --- | --- | --- |
| Human Piezo1 | ATCGCCATCATCTGGTTCCC | TGGTGAACAGCGGCTCATAG | NM_001142864.4 |
| Human GLP-1 | GCACATTCACCAGTGACTACAGCA | TGGCAGCTTGGCCTTCCAAATA | NM_002054.5 |
| Human β-actin | TCATGAAGATCCTCACCGAG | CATCTCTTGCTCGAAGTCCA | NM_001101.5 |
| Mouse Piezo1 | GCAGTGGCAGTGAGGAGATT | GATATGCAGGCGCCTATCCA | XM_011248372.2 |
| Mouse GLP-1 | ATTGCCAAACGTCATGATGA | GGCGACTTCTTCTGGGAAGT | NM_008100.4 |
| Mouse CCK | TAGCGCGATACATCCAGCAGGT | GGTATTCGTAGTCCTCGGCACT | NM_031161.5 |
| Mouse β-actin | CCACAGCTGAGAGGGAAATC | AAGGAAGGCTGGAAAAGAGC | NM_007393.5 |

**Supplementary information, Table S3. Key resources table**

| Reagent type (species) or resource  resource  resource | Designation | Source or reference | Identifiers | Additional  information |
| --- | --- | --- | --- | --- |
| Antibody | Mouse anti-GLP-1 | Abcam | Cat# ab23468, RRID:AB_470325 | WB: 1:1000  IF: 1:500 |
| Antibody | Rabbit anti-Piezo1 | Affinity Biosciences | Cat# DF12083, RRID:AB_2844888 | WB: 1:1000  IF: 1:400 |
| Antibody | Rabbit anti-Phospho-CaMK4 (Thr200) | Affinity Biosciences | Cat# AF3460, RRID:AB_2834898 | WB: 1:1000 |
| Antibody | Rabbit anti-CaMKIV | Cell Signaling Technology | Cat# 4032, RRID:AB_2068389 | WB: 1:1000 |
| Antibody | Rabbit anti-Phospho-mTOR (Ser2448) | Cell Signaling Technology | Cat# 5536, RRID:AB_10691552 | WB: 1:1000 |
| Antibody | Rabbit anti-mTOR | Cell Signaling Technology | Cat# 2983, RRID:AB_2105622 | WB: 1:1000 |
| Antibody | Rabbit anti-phospho-p70 S6 Kinase (Thr389) | Cell Signaling Technology | Cat# 9234, RRID:AB_2269803 | WB: 1:1000 |
| Antibody | Rabbit anti-p70 S6 Kinase | Cell Signaling Technology | Cat# 2903, RRID:AB_1196657 | WB: 1:1000 |
| Antibody | Rabbit anti-phospho-S6 Ribosomal Protein (Ser235/236) | Cell Signaling Technology | Cat# 4858, RRID:AB_916156 | WB: 1:1000 |
| Antibody | Rabbit anti-S6 Ribosomal Protein | Cell Signaling Technology | Cat# 2217, RRID:AB_331355 | WB: 1:1000 |
| Antibody | Mouse anti-β-actin | Cell Signaling Technology | Cat# 3700, RRID:AB_2242334 | WB: 1:1000 |
| Antibody | Goat anti-mouse fluorescein isothiocyanate-conjugated IgG | EarthOx LLC | Cat# E031210-01 |  |
| Antibody | Dylight 594 affinipure donkey anti-rabbit IgG | EarthOx LLC | Cat# E032421-01 |  |
| Antibody | Horseradish peroxidase‐conjugated, Goat Anti-Rabbit IgG | Jackson ImmunoResearch Labs | Cat# 111-035-003, RRID:AB_2313567 |  |
| Antibody | Horseradish peroxidase‐conjugated, Goat Anti-Mouse IgG | Jackson ImmunoResearch Labs | Cat# 115-035-003, RRID:AB_10015289 |  |
| Antibody | Mouse anti-CaMKKβ | Santa Cruz Biotechnology | Cat# sc-271674, RRID:AB_10708844 | WB: 1:1000 |
| Reagent | 0.1% gelatine | Biological Industries | Cat# 01-944-1B |  |
| Reagent | DMEM high sugar medium | Gibco | Cat# 11965092 |  |
| Reagent | Fetal bovine serum | Gibco | Cat# 12484028 |  |
| Reagent | Equine serum | Gibco | Cat# 16050122 |  |
| Reagent | Immobilon western chemiluminescent HRP substrate | Millipore | Cat# WBKLS0500 |  |
| Reagent | Diprotin A | Sigma-Aldrich | Cat# 90614-48-5 |  |
| Reagent | Thermo Scientific TurboFect Transfection Reagent | Thermo Fisher Scientific | Cat# R0531 |  |
| Reagent | TRIzol | Thermo Fisher Scientific | Cat# 15596026 |  |
| Reagent | RIPA Lysis Buffer | Beyotime Biotechnology | Cat# P0013B |  |
| Reagent | GsMTx4 | Alomone Labs | Cat# STG-100 |  |
| Reagent | Rapamycin | Santa Cruz Biotechnology | Cat# sc-3504B |  |
| Reagent | STO-609 | Selleck | Cat# S8274 |  |
| Reagent | Yoda1 | Sigma-Aldrich | Cat# SML1558 |  |
| Reagent | Dimethyl sulfoxide | Sigma-Aldrich | Cat# D2650 |  |
| Reagent | Exendin-4 | Sigma-Aldrich | Cat# E7144 |  |
| Reagent | Fluo-4 AM | Thermo Fisher Scientific | Cat# F14201 |  |
| Kit | Mouse Glucagon-Like Peptide 1 (GLP-1) ELISA Kit | Millipore | Cat# EGLP-35K |  |
| Kit | RT-PCR kit | Takara | Cat# RR014A |  |
| Oligonucleotides | Primers for cloning and RT-PCR, see Tables S1 and S2 | This paper |  |  |
| Fodder | Normal chow diet | Research Diets | Cat# D12450B |  |
| Fodder | High fat diet | Research Diets | Cat# D12492 |  |
